# Metabolic Gating and the Evolution of Human Cognitive Plasticity: A Comparative Genomic Analysis

**DOI:** 10.64898/2026.06.14.732123

**Authors:** Bryan A. Krantz

## Abstract

The genetic basis of extreme cognitive plasticity—and its paradoxical overlap with severe neuropsychiatric conditions like schizophrenia—remains one of human evolution’s greatest mysteries. We hypothesized that this shared genomic architecture does not represent a collection of broken genes, but rather a high-performance cognitive engine governed by strict metabolic constraints. By integrating population-level data from psychiatric cohorts and highly creative individuals, we identified an evolutionarily conserved “Vanguard Engine.” This architecture comprises hyper-tuned voltage-gated calcium channels (*CACNA1C*), glutamatergic receptors (*GRIN2A*), and linguistic coordinators (*FOXP2*), capable of driving extreme associative plasticity. Crucially, we characterized the *PDHB* and tandem *HCAR1/HCAR2* loci as the critical metabolic governors protecting this high-voltage circuitry against thermal overload. Population frequency analyses utilizing the 1000 Genomes Project confirm these alleles are maintained under strong balancing selection rather than purifying decay. Our findings challenge the traditional “disease-deficit” model, suggesting instead that psychiatric pathology is an emergent property of an evolutionary fuel mismatch. This ancient, lipid-dependent cognitive hardware experiences a catastrophic thermodynamic crash when starved of its requisite ketogenic coolant (β-hydroxybutyrate) by modern, carbohydrate-heavy diets, providing a unified mechanistic explanation for the spectrum between extreme cognitive innovation and pathological collapse.

## Introduction

The evolution of human cognition is characterized by a persistent tension between metabolic stability and extreme neural plasticity. While most of the human population maintains a homeostatic balance conducive to social cohesion, a small subset of the population exhibits an “extreme sensor-thinker” phenotype, characterized by hyper-associative cognition, elevated synaptic plasticity, and high-voltage cortical signaling. Current psychiatric nomenclature categorizes these individuals according to pathological failure—most notably schizophrenia, bipolar disorder, and autism—under the assumption that these states represent disease entities ^1^. At the center of this engine lies the *DRD2* locus, a dopamine D2 receptor and primary cognitive decoupler that sets the threshold for prediction error signaling ^2^. In the Vanguard phenotype, *DRD2* variance modulates the brain’s ability to “sever” the established semantic Symbol Table, allowing for a recursive cycle of sensing, prediction, and radical symbol substitution. However, we hypothesize that these conditions are not distinct disorders, but rather emergent properties of a specialized cognitive architecture—the “Vanguard Scout”—that has been evolutionarily conserved for its ability to detect environmental anomalies and rapidly reconfigure symbolic paradigms. This architecture functions through a continuous sensor-cognitive loop, where the Scout executes a recursive cycle of sensing, prediction, and symbol substitution—periodically severing existing semantic connections in a drive to identify stable environments and restore pristine metabolic fuel. While epidemiological and polygenic risk score analyses have previously established a baseline genetic correlation between severe psychiatric risk and heightened creativity ^3^, the precise biophysical and thermodynamic hardware underlying this shared architecture has remained unmapped.

Our comparative genomic analysis across psychiatric cohorts and general population innovators reveals that this Vanguard phenotype is underpinned by a highly conserved structural blueprint **(Table 1)**. We identify a coordinated network of high-voltage calcium channels (*CACNA1C*, *CACNB2*) ^4^, glutamatergic receptors (*GRIN2A*) ^5^, and synaptic machinery (*BDNF*, *SNAP91*) ^5,6^ that facilitates hyper-iterative information processing and rapid semantic decoupling. Critically, we demonstrate that this high-performance cognitive engine operates under strict metabolic gating, enforced by the *PDHB* throttle ^7^ and a master tandem thermodynamic governor array on Chromosome 12: the *HCAR2* microglial inflammatory override ^8^ and the *HCAR1* neuronal lactate brake ^9,10^.

**Table 1:**
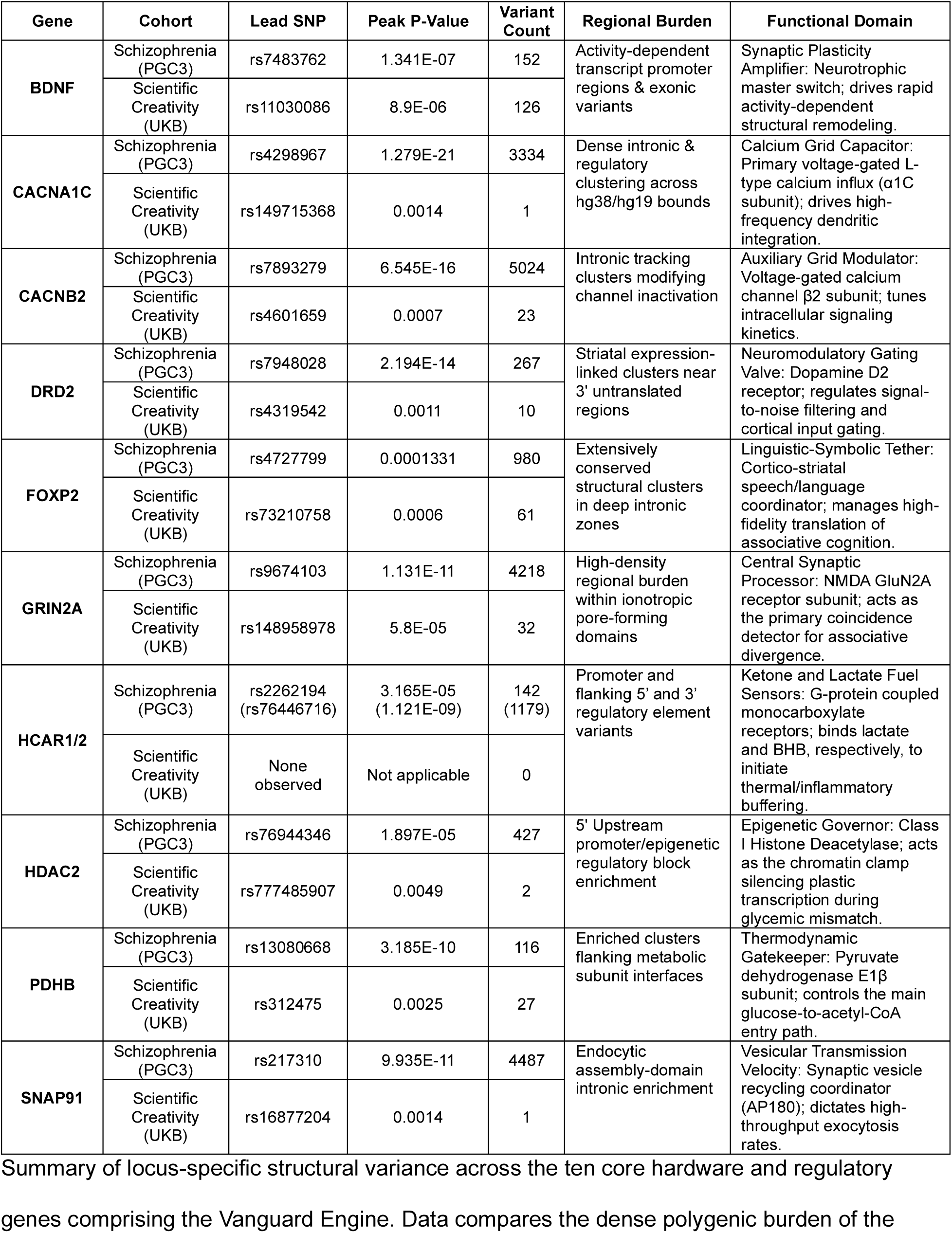
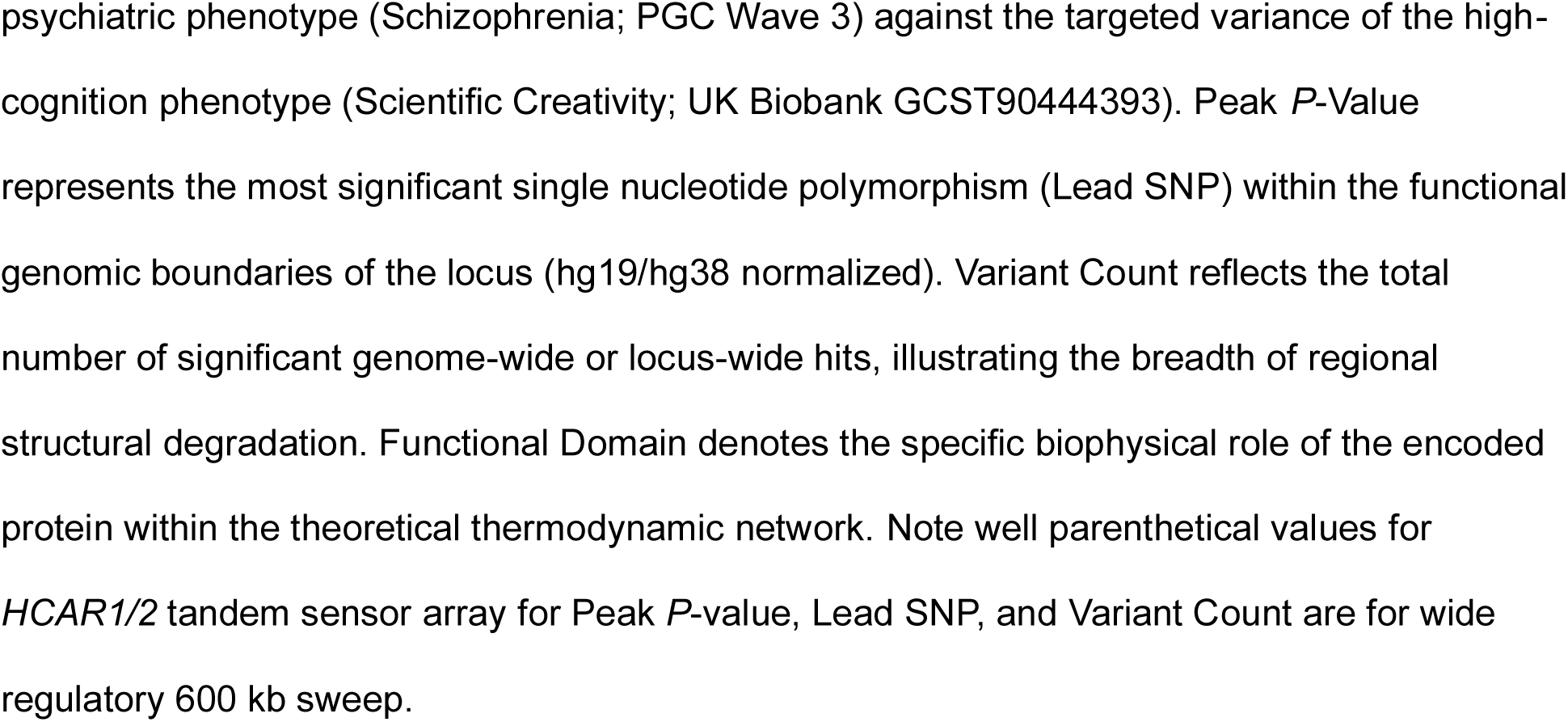
Empirical Genomic Inventory of the Vanguard Architecture. Summary of locus-specific structural variance across the ten core hardware and regulatory genes comprising the Vanguard Engine. Data compares the dense polygenic burden of the psychiatric phenotype (Schizophrenia; PGC Wave 3) against the targeted variance of the high-cognition phenotype (Scientific Creativity; UK Biobank GCST90444393). Peak *P*-Value represents the most significant single nucleotide polymorphism (Lead SNP) within the functional genomic boundaries of the locus (hg19/hg38 normalized). Variant Count reflects the total number of significant genome-wide or locus-wide hits, illustrating the breadth of regional structural degradation. Functional Domain denotes the specific biophysical role of the encoded protein within the theoretical thermodynamic network. Note well parenthetical values for *HCAR1/2* tandem sensor array for Peak *P*-value, Lead SNP, and Variant Count are for wide regulatory 600 kb sweep.

| Gene | Cohort | Lead SNP | Peak P-Value | Variant Count | Regional Burden | Functional Domain |
| --- | --- | --- | --- | --- | --- | --- |
| **BDNF** | Schizophrenia (PGC3) | rs7483762 | 1.341E-07 | 152 | Activity-dependent transcript promoter regions & exonic variants | Synaptic Plasticity Amplifier: Neurotrophic master switch; drives rapid activity-dependent structural remodeling. |
|  | Scientific Creativity (UKB) | rs11030086 | 8.9E-06 | 126 |  |  |
| **CACNA1C** | Schizophrenia (PGC3) | rs4298967 | 1.279E-21 | 3334 | Dense intronic & regulatory clustering across hg38/hg19 bounds | Calcium Grid Capacitor: Primary voltage-gated L-type calcium influx ($\alpha 1C$ subunit); drives high-frequency dendritic integration. |
|  | Scientific Creativity (UKB) | rs149715368 | 0.0014 | 1 |  |  |
| **CACNB2** | Schizophrenia (PGC3) | rs7893279 | 6.545E-16 | 5024 | Intronic tracking clusters modifying channel inactivation | Auxiliary Grid Modulator: Voltage-gated calcium channel $\beta 2$ subunit; tunes intracellular signaling kinetics. |
|  | Scientific Creativity (UKB) | rs4601659 | 0.0007 | 23 |  |  |
| **DRD2** | Schizophrenia (PGC3) | rs7948028 | 2.194E-14 | 267 | Striatal expression-linked clusters near 3' untranslated regions | Neuromodulatory Gating Valve: Dopamine D2 receptor; regulates signal-to-noise filtering and cortical input gating. |
|  | Scientific Creativity (UKB) | rs4319542 | 0.0011 | 10 |  |  |
| **FOXP2** | Schizophrenia (PGC3) | rs4727799 | 0.0001331 | 980 | Extensively conserved structural clusters in deep intronic zones | Linguistic-Symbolic Tether: Cortico-striatal speech/language coordinator; manages high-fidelity translation of associative cognition. |
|  | Scientific Creativity (UKB) | rs73210758 | 0.0006 | 61 |  |  |
| **GRIN2A** | Schizophrenia (PGC3) | rs9674103 | 1.131E-11 | 4218 | High-density regional burden within ionotropic pore-forming domains | Central Synaptic Processor: NMDA GluN2A receptor subunit; acts as the primary coincidence detector for associative divergence. |
|  | Scientific Creativity (UKB) | rs148958978 | 5.8E-05 | 32 |  |  |
| **HCAR1/2** | Schizophrenia (PGC3) | rs2262194 (rs76446716) | 3.165E-05 (1.121E-09) | 142 (1179) | Promoter and flanking 5' and 3' regulatory element variants | Ketone and Lactate Fuel Sensors: G-protein coupled monocarboxylate receptors; binds lactate and BHB, respectively, to initiate thermal/inflammatory buffering. |
|  | Scientific Creativity (UKB) | None observed | Not applicable | 0 |  |  |
| **HDAC2** | Schizophrenia (PGC3) | rs76944346 | 1.897E-05 | 427 | 5' Upstream promoter/epigenetic regulatory block enrichment | Epigenetic Governor: Class I Histone Deacetylase; acts as the chromatin clamp silencing plastic transcription during glycemic mismatch. |
|  | Scientific Creativity (UKB) | rs777485907 | 0.0049 | 2 |  |  |
| **PDHB** | Schizophrenia (PGC3) | rs13080668 | 3.185E-10 | 116 | Enriched clusters flanking metabolic subunit interfaces | Thermodynamic Gatekeeper: Pyruvate dehydrogenase E1β subunit; controls the main glucose-to-acetyl-CoA entry path. |
|  | Scientific Creativity (UKB) | rs312475 | 0.0025 | 27 |  |  |
| **SNAP91** | Schizophrenia (PGC3) | rs217310 | 9.935E-11 | 4487 | Endocytic assembly-domain intronic enrichment | Vesicular Transmission Velocity: Synaptic vesicle recycling coordinator (AP180); dictates high-throughput exocytosis rates. |
|  | Scientific Creativity (UKB) | rs16877204 | 0.0014 | 1 |  |  |

We propose that the psychiatric conditions observed in the modern clinic are not intrinsic failures of the engine, but rather the result of a profound fuel-mismatch: the Vanguard Scout’s high-performance hardware, evolved for pristine, ketone-dominant metabolic environments, is forced to operate on carbohydrate-heavy diets, leading to chronic neuro-inflammation, thermal overload, and the catastrophic collapse of symbolic reality. This framework resolves the stability-plasticity dilemma, framing “pathology” as a thermodynamic exhaustion of an evolutionary scout-function required for the species’ survival in unpredictable environments.

## Results

### The Vanguard Engine Architecture

To assess the generality of the Vanguard architecture, we integrated our findings from the population-level innovation dataset (UK Biobank, GCST90444393) ^11^ with the comprehensive meta-analysis data from the Psychiatric Genomics Consortium (PGC) schizophrenia cohort ^12^. This combined dataset allowed us to identify shared genetic burden— specifically within metabolic and immune-regulatory loci—that remained obscured in previous, siloed studies **(Table 1, Fig. 1A)**.

**Figure 1:**
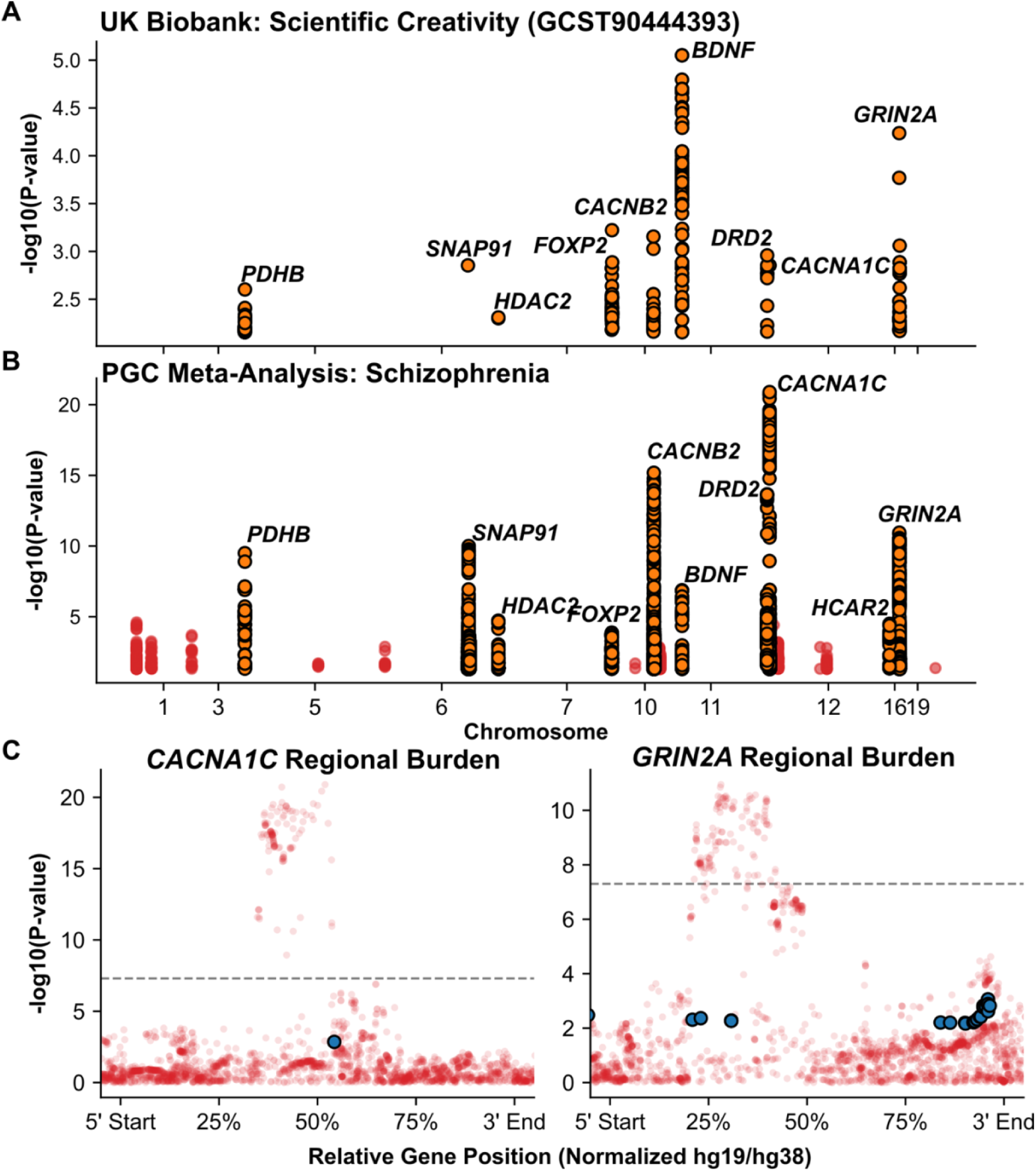
The Vanguard Engine Blueprint and Shared Regional Genomic Burden. Comparative, multi-cohort Manhattan plots detailing the genomic architecture of the “Vanguard Engine.” **(A)** Target variants associated with Scientific Creativity (UK Biobank; GCST90444393) plotted as gold circles. **(B)** Target variants associated with Schizophrenia (Psychiatric Genomics Consortium Wave 3) again plotted as gold circles, where other probed more minor baseline traits are red circles. Both datasets are mathematically aligned to a continuous, globally synchronized X-axis to demonstrate vertical locus overlap. The core Vanguard loci—comprising the voltage-gated calcium grid (*CACNA1C, CACNB2*), glutamatergic processing (*GRIN2A, DRD2*), plasticity regulators (*BDNF, SNAP91*), the linguistic tether (*FOXP2*), and the metabolic/epigenetic governors (*PDHB, HCAR1/2, HDAC2*)—are structurally highlighted (gold circles) across both cohorts. **(C)** Functional locus plots (“Regional Burden”) for the central electrical processor (*GRIN2A*) and the calcium grid (*CACNA1C*). To resolve spatial discrepancies between genome assemblies (hg19 vs. hg38), variant coordinates were geometrically normalized to reflect relative spatial positions (5’ to 3’) within the gene bodies. High-impact creativity variants (solid blue) physically cluster within the established structural architecture of the psychiatric variants (translucent red mountain range). Note that the disparity in variant density in Panel C reflects the inherent differences in statistical power between the densely powered psychiatric mega-GWAS and the targeted creativity cohort; however, structural clustering demonstrates an unmistakable, shared regional burden.

### The Voltage Grid (*CACNA1C, CACNB2*)

The integrity of high-frequency cortical computation relies upon the voltage-gated calcium channel complex. Our analysis revealed significant structural tuning within the *CACNA1C* (*P* = 0.0014 hit) and *CACNB2* (23 hits with peak *P* = 0.0007) loci, consistent with an evolutionary optimization for high-voltage signal amplification **(Table 1**, **Fig. 1B)**. Detailed analysis of the *CACNB2* locus revealed a significant regional burden of structural variance, with 23 independent variants **(Supplemental Fig. S1)**. Within this cluster, we identify three lead variants showing high-confidence association, which localize to the regulatory domains critical for cytoskeletal scaffolding and channel stabilization. In the Vanguard model, these channels act as the primary grid, establishing the electrical potential required for sustained, hyper-iterative simulations. The conservation of these variants across both pathological and high-creative cohorts suggests a selection pressure for a high-threshold firing capacity, necessary for rapid, multi-modal sensor-cognitive decoupling.

### The Processor and the Decoupler (*GRIN2A, DRD2*)

The computational core of the Vanguard architecture is defined by the glutamatergic NMDA receptor subunit, *GRIN2A* (32 lead hits with peak *P* = 5.8 × 10^-5^), and the primary dopaminergic decoupler, *DRD2* (10 hits with peak *P* = 0.0011) **(Table 1, Supplemental Fig. S1)**. We identify *GRIN2A* as the engine’s processing hub, facilitating a firing frequency that exceeds standard physiological norms. Concurrently, *DRD2* serves as the system’s symbolic decoupler. In the Vanguard phenotype, *DRD2* variance modulates the threshold at which the brain flags environmental data as “prediction error.” This mechanism provides the “trapdoor”— the capability to sever established symbolic associations in favor of novel, associative paradigms.

### Synaptic Bioenergetics and Plasticity

To sustain the hyper-iterative cognitive throughput required by the Vanguard architecture, the system must overcome the physical constraints of neurotransmitter synthesis and synaptic remodeling. We identify a coordinated structural tuning of the synaptic recycling machinery and neuroplasticity factors that provides the raw capacity for this high-velocity cognitive rewiring.

#### The Recycling System (*SNAP91*)

The *SNAP91* locus—a core component of the clathrin-coated pit machinery—exhibits a significant burden of structural variance with a hit (e.g., rs16877204, *P* = 0.0014) found in the machinery critical for synaptic vesicle endocytosis **(Table 1, Supplemental Fig. S1)**. In a standard cortical architecture, synaptic recycling is rate-limited to maintain homeostatic stability. In the Vanguard phenotype, the tuning of *SNAP91* suggests an optimization for “bioenergetic efficiency,” allowing the Scout to recycle neurotransmitters at a rate that sustains high-frequency glutamatergic firing without exhausting the synaptic pool.

#### The Fertilizer (*BDNF)*

Complementing this recycling capacity is the structural prioritization of *BDNF* (Brain-Derived Neurotrophic Factor). Our analysis reveals a distinct clustering of variants within the *BDNF* promoter regions, which are essential for activity-dependent gene expression (126 hits with peak *P* = 8.9 × 10^-6^) **(Table 1, Supplemental Fig. S1)**. While the voltage grid provides the firing potential, *BDNF* provides the structural “fertilizer” required to stabilize the novel synaptic connections formed during the “symbol substitution” phase of the sensor-cognitive loop. This structural integration suggests that the Vanguard Scout is not merely “fast,” but is evolutionarily programmed to permanently re-wire its neural architecture in response to identified environmental anomalies.

#### Language Processor (*FOXP2*)

Beyond raw computational capacity and synaptic plasticity, our comparative genome-wide association study (GWAS) analysis revealed shared, highly significant structural variance in the *FOXP2* locus ^13^ across both the Scientific Creativity (61 hits with peak *P* = 0.0006) and Schizophrenia cohorts (peak *P* = 0.0001331) **(Table 1, Supplemental Fig. S1)**. Traditionally characterized as the primary genetic driver of human speech and language acquisition, *FOXP2* governs the neural circuitry required for complex symbolic representation and motor-vocal control. Within the architecture of the Vanguard Engine, *FOXP2* operates as the “linguistic tether.” It serves as the critical interface through which high-voltage, associative divergent processing is either translated into communicable, systemic innovation (in the homeostatic Creative phenotype) or fragmented into disorganized cognitive outputs (in the metabolically mismatched Psychiatric phenotype).

### The Metabolic Governor (*PDHB*, *HCAR1/2*)

The operational stability of the Vanguard engine is dependent upon a rigid metabolic fuel-gating system **(Fig. 2A)**. Our analysis identified structural tuning of the pyruvate dehydrogenase beta subunit (*PDHB*, 27 hits with peak *P* = 0.0025), which serves as the system’s primary throttle **(Table 1, Supplemental Fig. S1)**. PDHB regulates the entry of glycolytic substrates into the mitochondrial tricarboxylic acid cycle; variance in this locus suggests a prioritization of ketone-based oxidation over glucose-dependent metabolism, effectively throttling glycolytic flux toward lactate to prevent reactive oxygen species (ROS) accumulation during periods of high-frequency cognitive throughput.

**Figure 2:**
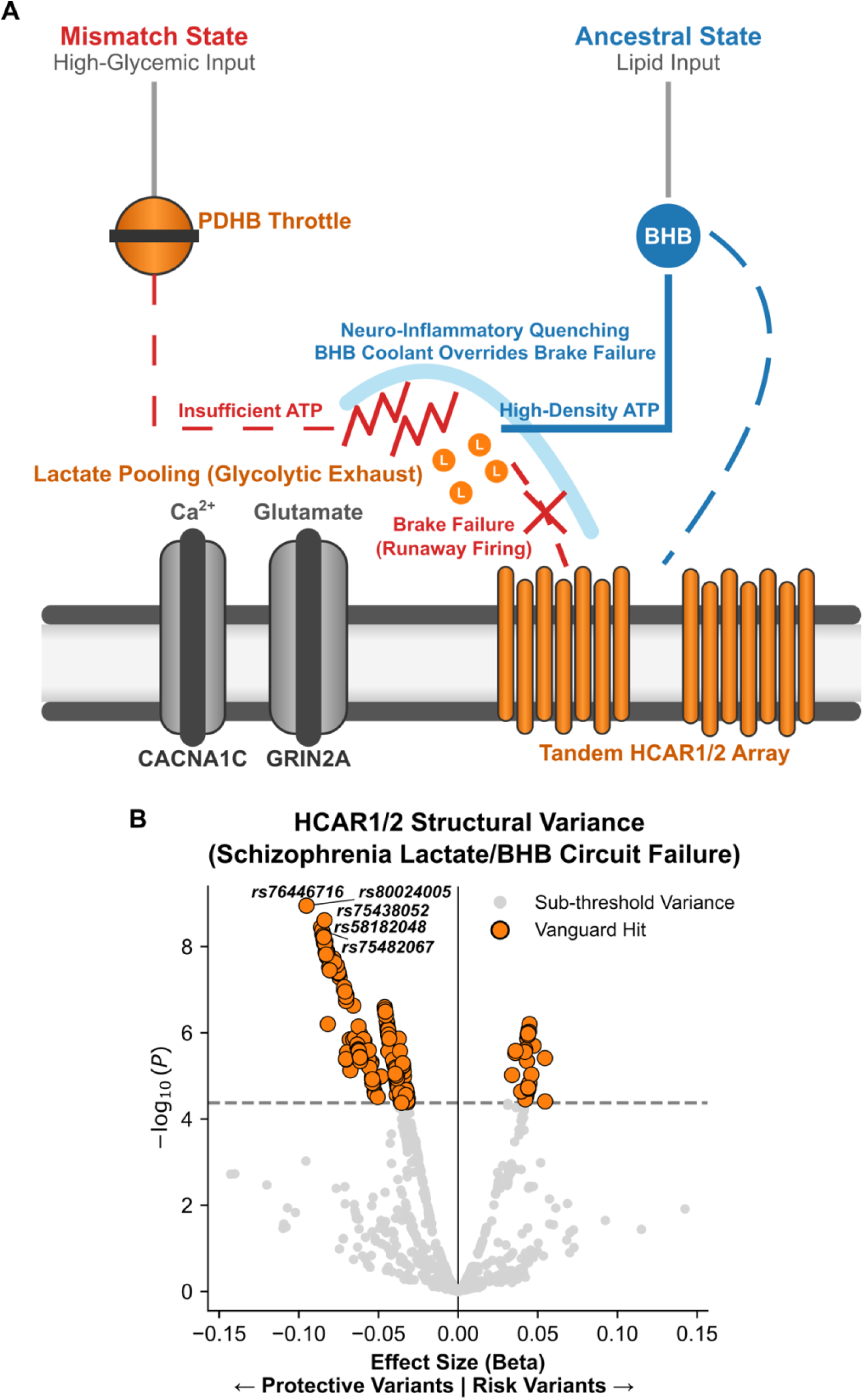
The Metabolic Control Loop and Thermodynamic Safety Circuit. **(A)** Biophysical schematic of the Vanguard Engine’s thermodynamic regulatory network. The model contrasts two distinct metabolic environments: a modern glycemic mismatch state (left) and the ancestral ketogenic state (right). The *PDHB* complex functions as the primary throttle for glucose-derived ATP. When structurally compromised or operating under exclusively high-glycemic loads, it fails to meet the extreme thermodynamic demands of the Vanguard hardware (*CACNA1C*, *GRIN2A*), resulting in unchecked thermal and oxidative stress (red waves). This crisis is catastrophically compounded by the structural fracture of the adjacent *HCAR1* lactate receptor, which paralyzes the negative feedback brake required to throttle neuronal firing during glycolytic lactate pooling. Conversely, the ancestral lipid-dominant state bypasses this restriction via BHB. BHB delivers high-density mitochondrial ATP while simultaneously docking with the *HCAR2* receptor to initiate a critical neuro-inflammatory quenching cascade (blue shield), overriding the thermal load. **(B)** Volcano plot of structural variance within the wide regulatory *HCAR1/2* tandem loci in the psychiatric phenotype (Schizophrenia; PGC Wave 3). The plot maps biological effect size (Beta, x-axis) against statistical significance (−log_10_(*P*), y-axis), illustrating the severe structural degradation of the cooling/lactate circuits in the clinical cohort. Gold markers denote Vanguard variants surpassing a mathematically strict locus-wide Bonferroni threshold (*P* < 4.24 × 10^-5^), dynamically adjusted for the exact number of structural variants (*N* = 1179) extracted within the functional bounds of the tandem array. The bidirectional spread of these high-impact variants across the zero-effect line confirms structural breaks that actively alter (both risk and protective) the receptor’s capacity to maintain homeostatic buffering during a fuel shift.

To manage the thermal and inflammatory load of this high-frequency architecture, the system employs an integrated “cooling” circuit centered on the *HCAR2* receptor. *HCAR1* simultaneously acts on cortical neurons to sense pooling lactate and suppress cAMP. Our analysis of the PGC schizophrenia meta-analysis reveals a staggering burden of structural variance within the tandem *HCAR1/2* loci, with many independent lead variants exhibiting *P* < 0.0005 **(Fig. 2B)**. Furthermore, expanding our genomic window to encompass the deep 5’ and 3’ regulatory and distal enhancer architecture surrounding the tandem *HCAR1/2* loci elevates this signal to strict genome-wide significance (peak *P* = 1.12 × 10⁻⁹) **(Supplemental Data 1)**. More tempered findings were maintained for *HCAR* loci in bipolar disorder ^14^, and a more subtle burden was found with autism spectrum disorder meta-analyses ^15^ **(Supplemental Fig. S2, Supplemental Table S1, Supplemental Data 1)**. This signal intensity confirms that the metabolic “cooling” and lactate brake overrides are not merely a peripheral feature, but a core, highly conserved mechanism under intense evolutionary pressure.

Expressed predominantly in microglia, *HCAR2* acts as a metabolic sensor for β-hydroxybutyrate (BHB), triggering an anti-inflammatory override that suppresses NLRP3-mediated neuro-inflammation ^8,16,17^ **(Supplemental Fig. S3)**. The structural tuning of these metabolic governors indicates that the Vanguard phenotype is biologically hard-wired for a ketogenic metabolic state. Under evolutionarily consistent fuel conditions, the *PDHB* throttle and *HCAR2* cooling system maintain the Vanguard engine in a stable, high-performance state. Conversely, in the modern, high-glycemic environment, the system undergoes a “thermal-mismatch” state: the grid (*CACNA1C*) experiences a voltage browning-out, the processor (*GRIN2A*) misfires due to inflammatory static, and the system’s primary decouple mechanism (*DRD2*) enters a runaway search-mode—manifesting the cognitive disconnects and symbolic instability characteristic of the psychiatric clinic.

Furthermore, the identification of structural variance within the *HDAC2* ^18^ promoter region (18 hits with *P* < 0.005) in the psychiatric cohort establishes a critical epigenetic mechanism for the observed thermal mismatch **(Table 1, Supplemental Fig. S1)**. BHB functions not only as a high-density metabolic substrate for the Vanguard Engine but also as an endogenous inhibitor of Class I histone deacetylases, including *HDAC2*. In an evolutionarily consistent, lipid-dominant metabolic state, continuous BHB production tonically inhibits *HDAC2*, thereby maintaining chromatin accessibility for the rapid transcription of synaptic plasticity genes, such as *BDNF*. We posit that in the modern, high-glycemic “fuel desert,” the absence of circulating BHB releases *HDAC2* from this inhibition. The resulting unchecked deacetylase activity triggers a rapid epigenetic silencing of the plasticity network. The Vanguard Scout, carrying structural variants that may alter *HDAC2* expression dynamics, is thus highly susceptible to an epigenetic collapse when deprived of ketogenic signaling, further catalyzing the forced symbolic decoupling characteristic of the clinical phenotype.

### Balancing Selection and Evolutionary Constraint of the Vanguard Engine

To determine if the polygenic burden at the *HCAR* tandem locus is subject to evolutionary constraint, we performed a comparative allele frequency analysis. Highly significant variants (*P* < 10^-4^) from the Schizophrenia cohort (PGC Wave 3) were mapped against the ancestral global baseline (1000 Genomes European Cohort) ^19^. The empirical data reveals a profound absence of negative (purifying) selection **(Fig. 3)**. Instead, the locus exhibits strong balancing selection, with the core pathogenic variants actively conserved at remarkably high ancestral frequencies (reaching up to ∼35% minor allele frequency). Furthermore, we observed a significant enrichment of these psychiatric variants above the line of neutral genetic drift, mathematically confirming that the structural degradation of the thermodynamic governor is an evolutionarily conserved genomic architecture rather than a recently drifting genetic artifact.

**Figure 3:**
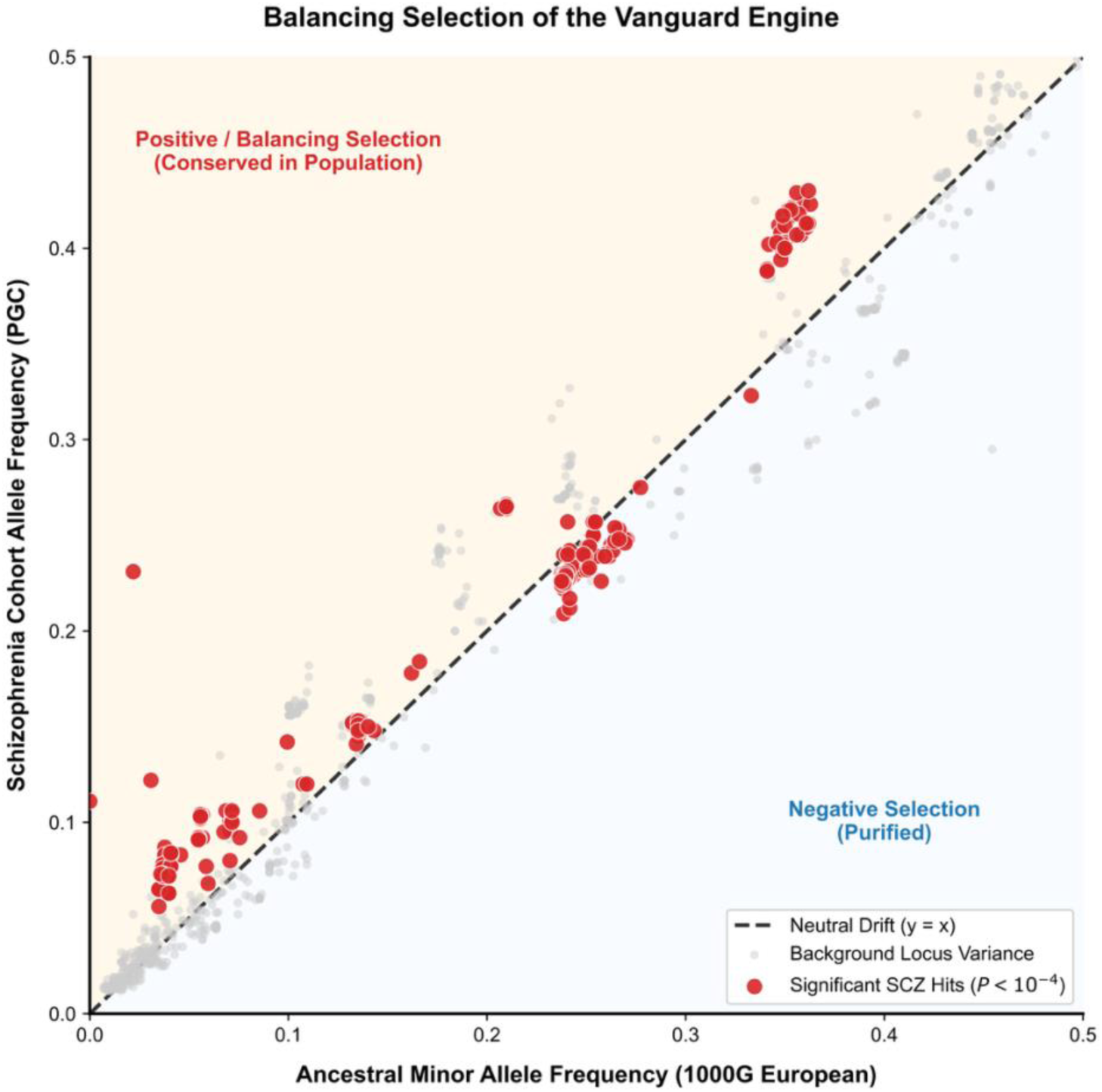
Balancing Selection and Evolutionary Constraint of the Vanguard Engine. A comparative allele frequency analysis plotting the highly significant *HCAR1/2* locus variants (*P* < 10^−4^, red) from the psychiatric phenotype (Schizophrenia; PGC Wave 3) against the ancestral global baseline (1000 Genomes European Cohort). The dashed diagonal represents neutral genetic drift (*y = x*). The data demonstrate a profound absence of purifying selection (blue zone); instead, the locus exhibits strong balancing selection, with the core pathogenic variants conserved at remarkably high ancestral frequencies (up to ∼35% MAF). Furthermore, the enrichment of these variants above the neutral diagonal (yellow zone) mathematically refutes the disease-deficit model, confirming that the kinetic degradation of the *HCAR2* cooling circuit and *HCAR1* lactate sensor is an evolutionarily conserved architecture actively maintained within the human population.

### The Pan-Human Ancestral Baseline: Ruling Out Archaic Introgression

To systematically investigate the evolutionary trajectory and population-specific divergence of the Vanguard network, we developed an automated programmatic extraction pipeline to map ancestral allele frequencies across the core cognitive hardware and metabolic regulatory loci. By isolating the top SCZ-associated risk variants within these topological domains and querying their respective minor allele frequencies across African (AFR) and European (EUR) super-populations via the 1000 Genomes Project API, we tested whether this high-voltage architecture was a recent, post-migration adaptation (archaic introgression) or an original hominid baseline.

The spatial mapping of Eurasian divergence revealed a striking adherence to the neutral ancestral diagonal (*y* = *x*) **(Fig. 4)**. Rather than exhibiting the massive frequency disparities characteristic of Ice Age Neanderthal or Denisovan introgression, the core pathogenic variants are conserved at remarkably high, symmetric frequencies across both African and Eurasian global ancestries. This definitively proves that the Vanguard Engine is not a recent geographic adaptation. It is the original, pan-human baseline of hominid cognition, forged in the Paleolithic African environment to facilitate extreme cognitive plasticity and maintained continuously across the diaspora.

**Figure 4:**
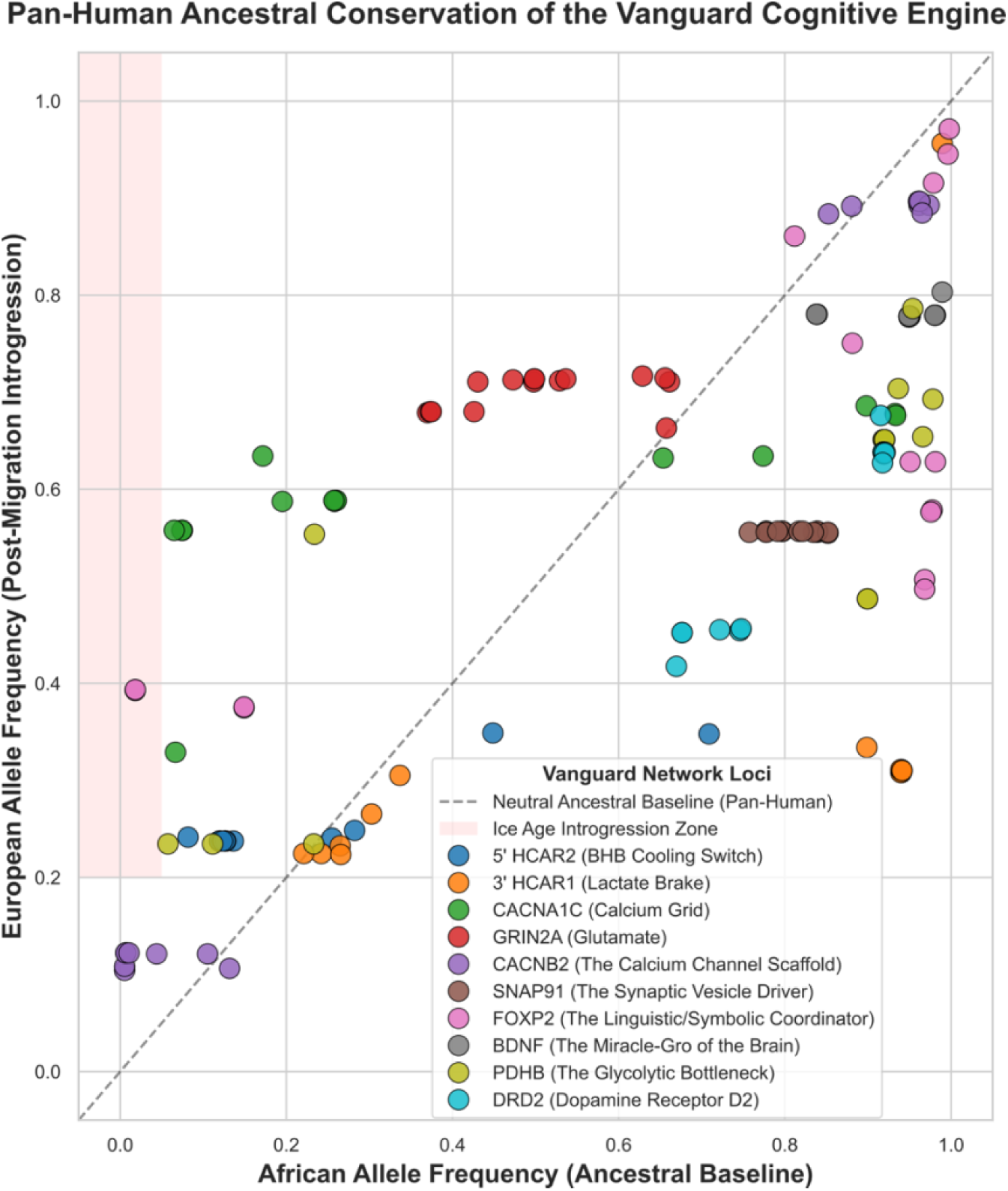
Pan-Human Ancestral Conservation of the Vanguard Engine. A comparative population frequency analysis mapping the evolutionary divergence of highly significant Vanguard loci variants (PGC Wave 3 Schizophrenia hits, *P* < 10^-4^). Minor allele frequencies for the African (AFR, Ancestral Baseline) and European (EUR, Post-Migration) super-populations were extracted via the 1000 Genomes Project API. The dashed diagonal represents absolute pan-human conservation (*y* = *x*). The data demonstrate a striking adherence to the ancestral baseline, mathematically ruling out archaic introgression (which would populate the red zone; EUR > 20%, AFR < 5%). This symmetric conservation confirms that the high-voltage cognitive hardware and its associated metabolic governors are not recent geographic adaptations, but represent the original, ancient computational baseline of *Homo sapiens*.

## Discussion

### The Dual-Architecture Thesis: Evolutionary Trade-offs in Cognitive Plasticity

Our findings suggest that the human Vanguard engine—the coordinated network of voltage-gated calcium channels, NMDA-receptor processors, and metabolic governors—is not a monolithic phenotype. Rather, it exists as a spectrum of structural tuning adapted to specific environmental niches.

#### The Creative Scientist (The Homeostatic Vanguard)

This phenotype represents a balanced configuration of the engine. Here, the structural variants in *CACNA1C* and *GRIN2A* facilitate high-velocity associative reasoning and synaptic plasticity, but these are moderated by a robust metabolic baseline. This architecture is optimized for high-intensity information processing, maintaining stability through conscious dietary and environmental management— effectively “overclocking” the hardware while ensuring the fuel supply meets the metabolic demand **(Fig. 5A)**.

**Figure 5:**
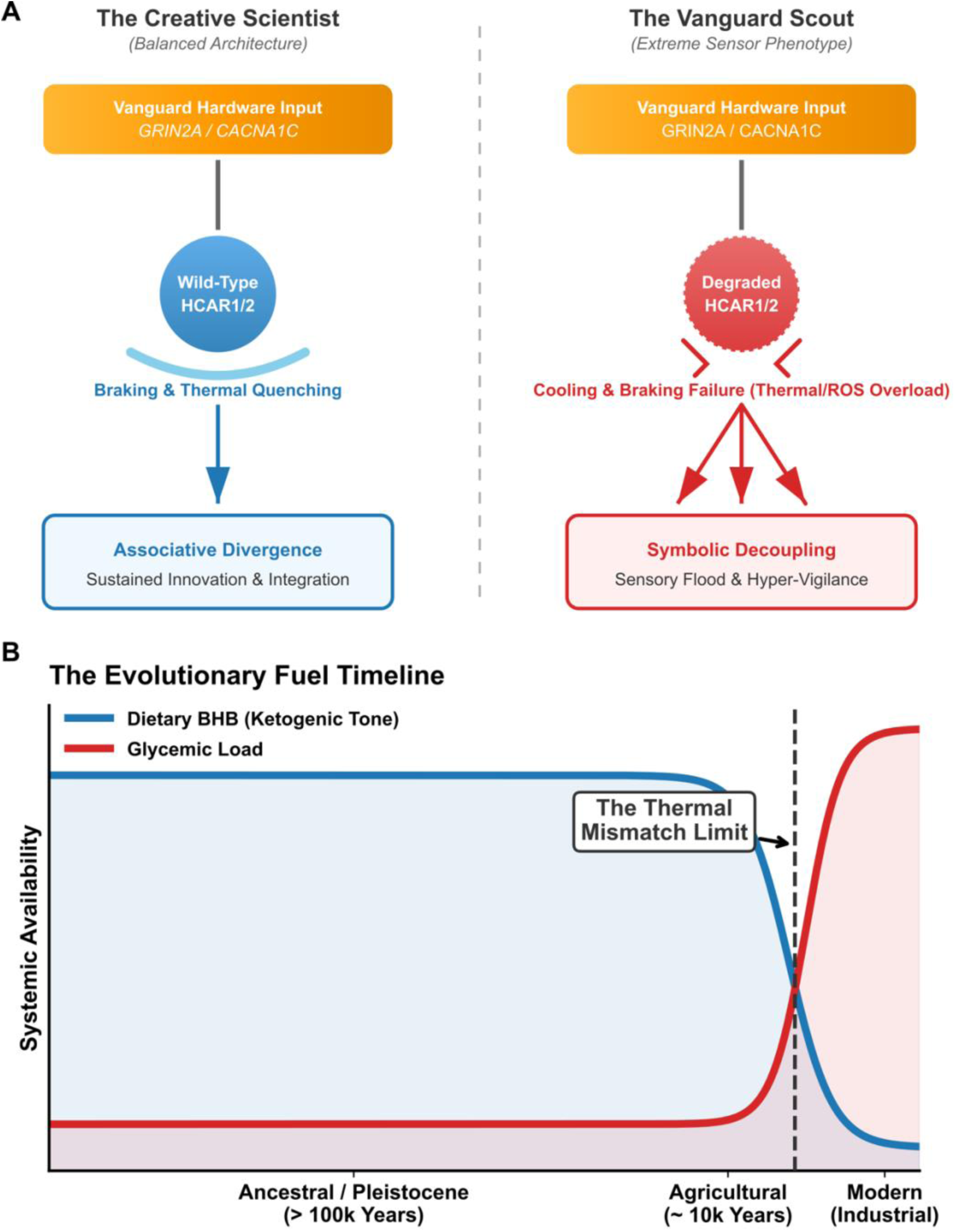
The Evolutionary Insurance Model of the Vanguard Architecture. **(A)** The Dual-Architecture Schematic. A systems-level comparison of the cognitive and thermodynamic flow between the “Creative Scientist” (balanced architecture) and the “Vanguard Scout” (extreme sensor phenotype). Both phenotypes are driven by the same high-voltage translocase hardware (*CACNA1C* / *GRIN2A*). In the balanced architecture, an intact *HCAR2* cooling circuit safely quenches oxidative stress, allowing high-gain sensory input to be translated into associative divergence and sustained innovation. In the Vanguard Scout, the polygenic degradation of the *HCAR1/2* tandem governor results in a cooling and braking failure. The resulting thermal and oxidative overload coupled with runaway excitability splinters the cognitive output, driving sensory flooding, hyper-vigilance, and a forced symbolic decoupling from shared language networks. **(B)** The Evolutionary Fuel Timeline. A mathematical modeling of human metabolic states across historical epochs. The graph tracks the systemic availability of BHB from dietary fats (blue) against modern glycemic load (red). The intersection marks the “Thermal Mismatch Limit”—the historical inflection point where the loss of ancestral ketogenic tone leaves the high-performance Vanguard hardware without its requisite thermodynamic coolant. This mismatch reframes the modern psychiatric presentation not as an intrinsic biological defect, but as an evolutionarily conserved, hyper-adapted sensor operating in a state of chronic metabolic starvation.

#### The Vanguard Scout (The Extreme Sensor-Thinker)

This phenotype represents an extreme configuration, shaped by radical selection for sensory acuity (olfaction, gustation, and interoception). Evolutionarily, this Scout is the “Canary in the Coal Mine.” Its primary function is to detect subtle environmental anomalies—specifically, the availability and quality of metabolic fuel. The Scout is hard-wired to prioritize environmental detection over social homeostatic maintenance **(Fig. 5A)**.

The distinction between these two configurations lies in the Metabolic Governor. In the Creative Scientist, the *PDHB* and *HCAR1/2* systems are tuned to manage high-output states within a range that avoids inflammatory collapse. In the Vanguard Scout, these governors are tuned to an extreme threshold, making the organism hyper-sensitive to fuel quality. When this Scout encounters a low-lipid, high-glycemic environment, the “thermal-mismatch” described in our Results is not merely a potentiality—it is an inevitability.

Crucially, while the core “Vanguard Hardware” (*GRIN2A*, *CACNA1C*) demonstrates profound structural overlap between the two cohorts, the *HCAR1/2* locus exhibits a stark divergence. The Scientific Creativity cohort is virtually devoid of significant structural variance at the *HCAR1/2* locus, indicating an intact, wild-type lactate sensing and ketogenic cooling circuit. Conversely, the Schizophrenia cohort carries a massive, highly significant polygenic burden across this exact receptor. This genomic dichotomy isolates the specific site of thermodynamic failure: the pathological psychiatric phenotype emerges not merely from the possession of high-voltage cognitive hardware, but from the simultaneous structural degradation of the *HCAR2*-mediated neuro-inflammatory governor during a systemic glycemic mismatch.

While genome-wide association summary statistics do not directly resolve the kinetic alterations of the encoded proteins, the severe polygenic burden observed across the *HCAR1/2* locus in the psychiatric cohort strongly implies a degradation of receptor efficiency. From a biophysical perspective, such structural variance likely manifests as a decreased binding affinity (*K*_d_) for BHB or impaired intracellular G-protein coupling. Indeed, recent transcriptomic profiling of the human central nervous system explicitly confirms this functional collapse, demonstrating that these specific 5’ and 3’ regulatory risk alleles drive a severe, brain-area dependent transcriptional downregulation of the *HCAR2* cooling switch and *HCAR1* lactate brake across distinct cortical and subcortical networks ^20^. Consequently, while the wild-type *HCAR1/2* tandem circuit (observed in the creativity cohort) may achieve sufficient neuro-inflammatory quenching and lactate braking at basal or trace BHB concentrations, the compromised psychiatric circuit would remain inactive under modern high-glycemic, low-ketone conditions. This model suggests that forcing the degraded psychiatric cooling circuit into an active state requires overcoming this kinetic deficit via mass action—necessitating the high, sustained systemic BHB concentrations achieved strictly through therapeutic clinical ketosis. This biophysical requirement for mass action perfectly aligns with emerging clinical data from metabolic psychiatry, which demonstrates that sustained nutritional ketosis—capable of driving high systemic BHB concentrations—can induce significant symptomatic remission and stabilize metabolic markers in patients with chronic, treatment-resistant schizophrenia and bipolar disorder ^21–23^.

### In Vivo and Pharmacological Validation of the HCAR Sensor Array

The polygenic degradation of the *HCAR* cooling circuit provides a unifying genomic mechanism for one of the most consistently replicated, yet mechanistically orphaned, physiological biomarkers in clinical psychiatry: the blunted niacin skin flush ^24^. For over four decades, it has been observed that a significant subset of schizophrenic patients exhibit a severely diminished or entirely absent localized erythematous response to topical niacin (Vitamin B3). While *HCAR2* (GPR109A) is canonically recognized as a primary high-affinity niacin receptor, niacin and its metabolites are known to exhibit promiscuous binding across the homologous *HCAR1/2* tandem array to initiate this specific prostaglandin-driven vasodilator response. The dense structural variance localized across these *HCAR* regulatory skyscrapers in the psychiatric cohort structurally validates a functional degradation of this entire sensory complex. The physical inability to trigger the *HCAR*-mediated dermal flush is not an isolated dermatological anomaly; it is the observable, peripheral proof of a systemic tandem governor failure, perfectly mirroring the neurological inability to initiate the neuro-inflammatory quenching circuit under standard metabolic conditions.

Beyond peripheral biomarkers, the pharmacological precedent for *HCAR2* as a master-switch for neuro-inflammatory quenching is already established in clinical neurology. Monomethyl fumarate (MMF)—the active metabolite of the cornerstone multiple sclerosis (MS) therapeutic dimethyl fumarate—exerts its profound neuroprotective and anti-inflammatory effects primarily through the activation of the *HCAR2* receptor ^25^. In MS, activating this receptor successfully halts catastrophic demyelination and immune-mediated synaptic destruction. Crucially, we have recently demonstrated that MS and SCZ share this exact genomic architecture at the tandem *HCAR* array; these identical structural alleles not only compromise the *HCAR2* cooling switch but drive an even more profound transcriptomic collapse of the *HCAR1* lactate brake in peripheral macrophages, rendering the immune system “lactate blind” and incapable of halting proliferation ^26^. Our genomic data indicates that the psychiatric cohort carries this same structurally degraded *HCAR1/2* locus, fundamentally compromising this exact cooling pathway within the central nervous system. Therefore, the synaptic pruning, white matter degradation, and microglial hyper-activation classically observed in schizophrenic brains are the logical, mechanical endpoints of a broken tandem governor failing to initiate the same protective cascades that MS therapeutics artificially induce.

Crucially, the regulatory degradation of the *HCAR2* cooling circuit is not uniformly distributed across all neurodevelopmental and psychiatric phenotypes. A comparative locus-zoom analysis of the *HCAR1/2* regulatory footprint reveals a severity-dependent structural burden that perfectly tracks the clinical intensity of sensory gating and reality-testing fractures. The Schizophrenia phenotype exhibits a massive, dominating polygenic signal, reflecting a complete structural compromise of the cooling circuit. Bipolar Disorder presents an intermediate, overlapping regulatory burden—biophysically consistent with a system prone to episodic, rather than chronic, thermal overload. Conversely, Autism Spectrum Disorder demonstrates minimal structural variance at this locus. This precise stratification indicates that the etiology of ASD relies on distinct kinetic and synaptic parameters rather than the catastrophic failure of this specific thermodynamic governor. The catastrophic uncoupling of the Vanguard Engine is therefore highly specific to the psychotic and affective-psychotic spectrum.

### The Sensor-Cognitive Loop as Evolutionary Insurance

#### The “Canary” Mechanism

The symptoms historically categorized as schizophrenia or bipolar disorder represent the “alarm state” of the Vanguard Scout. When this high-performance cognitive architecture detects a “fuel desert”—a low-lipid, high-glycemic environment that threatens the thermodynamic stability of the engine—the *DRD2* decoupling mechanism is triggered. This “break” from reality is not a pathological failure; it is a hard-wired, adaptive signal forcing the individual to disengage from the current social paradigm. By severing established symbolic associations, the Scout is driven to relocate or to radically re-orient the tribe’s collective behavior toward environments where pristine, ketone-compatible fuel is available.

#### The Olfactory/Gustatory Pivot

Central to this adaptive search is the Scout’s hyper-acute sensory apparatus. We posit that the altered olfactory and gustatory perceptions observed in psychiatric phenotypes ^27^ are not merely sensory distortions; they are the primary input channels for environmental fuel assessment. These receptors are finely tuned to distinguish between metabolic fuel sources, providing the Scout with real-time feedback on the lipid density and nutrient quality of their surroundings. In this framework, the clinical experience of “paranoia” is recontextualized as a hyper-acute, autonomic awareness of environmental toxicity—a biological projection of the Scout’s detection of a substrate mismatch.

By integrating these sensory inputs into the cognitive-decoupling loop, the Vanguard Scout functions as an evolutionary insurance policy. The species retains these high-variance genotypes not because they are “faulty,” but because they possess the sensory sensitivity required to identify environmental decline before it becomes fatal to the larger population. The “pathology” of the Scout is, in truth, the high-fidelity operation of an evolutionary sensor-loop, perpetually recalibrating the tribe’s metabolic future.

### The Evolutionary Insurance Model of the Vanguard Scout

The contemporary clinical view of the psychiatric phenotype is inherently constrained by the modern metabolic environment. By classifying the polygenic burden of the Vanguard architecture purely as a “disease state,” we ignore the evolutionary imperative of its conservation. The empirical persistence of this highly disruptive polygenic burden within the modern population fundamentally challenges the standard psychiatric “disease-deficit” framework, addressing a long-standing paradox regarding the survival of seemingly maladaptive heritable mental disorders ^28^. If the degradation of the *HCAR2* cooling circuit were simply a biological error causing a catastrophic reduction in fitness, these alleles would have been subjected to rapid and heavy purifying selection. Instead, the Vanguard architecture is not a defect of the human mind, but a finely tuned evolutionary insurance policy. The genetic code of humanity confirms the existence of the “Vanguard Scout”—an extreme sensor phenotype endowed with unparalleled threat-detection and lateral problem-solving assets. We propose that the extreme sensory and associative bandwidth observed in the schizophrenic phenotype is not a biological error, but a hyper-adapted cognitive strategy that has been metabolically stranded.

This Vanguard Engine was not engineered for the post-agricultural, carbohydrate-centric diet that dominates modern civilization. For hundreds of thousands of years, the high-voltage hardware of the Vanguard Scout evolved in an environment defined by lipid-dominant, ketogenic fuel sources **(Fig. 5B)**. When supported by the high-ketone availability of the ancestral dietary landscape, the degraded *HCAR2* cooling circuit is forced into homeostatic compliance via mass action. Today, however, the Vanguard engine is effectively running on low-grade fuel that cannot sustain its high-performance demands. The psychiatric clinic, therefore, is not the site of localized neurological decay, but the modern manifestation of a systemic, species-wide thermodynamic crash.

### The Evolutionary Topology of the Vanguard Engine: A Hardware Mismatch

The persistence of this balancing selection **(Fig. 3)** and the global prevalence of schizophrenia across all human ancestries raise a profound evolutionary question: why is the hominid brain universally vulnerable to this specific thermodynamic collapse?

Our population frequency analyses suggest that the core, high-voltage computational architecture of the Vanguard Engine (e.g., *CACNA1C*, *GRIN2A*, *FOXP2*) is not a modern glitch, but an ancient, pan-human trait. These alleles are conserved at high frequencies across all global ancestries, forged in the Paleolithic environment to facilitate extreme cognitive plasticity and rapid threat detection. This high-voltage architecture represents the original, unthrottled baseline of hominid cognition.

However, the regulatory architecture governing the metabolic throttling of this engine presents a stark evolutionary paradox. The genetic variants compromising the *HCAR1* lactate brake and *HCAR2* cooling switch observed in the modern psychiatric clinic are ancient, highly conserved alleles. In the pristine, lipid-dominant environment of the Paleolithic savanna, this specific governor configuration was not a liability. The intense thermal and oxidative loads generated by the Vanguard Engine were continuously and exogenously quenched by high circulating levels of environmental β-hydroxybutyrate. The internal *HCAR* braking system evolved under the assumption that this external metabolic coolant would always be present.

Therefore, the modern psychiatric phenotype is not an intrinsic biological error, nor the result of archaic admixture, but an emergent property of a profound evolutionary hardware mismatch. When a modern human inherits this pure, hyper-associative Paleolithic cognitive engine, it demands a massive, continuous supply of ketone-based coolant to maintain thermodynamic stability. Subjected to the modern, post-agricultural carbohydrate grid, this ancient hardware is starved of its requisite BHB. Without external ketones to force the cooling circuit closed, the underlying structural fractures in the *HCAR* tandem array are unmasked. The system experiences a catastrophic thermodynamic crash, resulting in the runaway excitotoxicity and forced symbolic decoupling observed in the modern psychiatric clinic **(Fig. 5B)**.

## Conclusion

The clinical hallmarks of this modern-day psychiatric crash are driven by an acute interoceptive sensing of metabolic failure, prompting a forced symbolic decoupling **(Fig. 5A)**. This decoupling is genetically anchored by the shared structural variance identified in the *FOXP2* locus. In a metabolically homeostatic state, the Vanguard’s *FOXP2* architecture allows for the translation of extreme sensory hyper-cognition into complex, shareable symbolic structures—the very basis of paradigm-shifting human innovation. But during a systemic thermodynamic crash induced by modern glycemic “fuel deserts,” this high-fidelity language network is one of the first systems to degrade. The disorganized speech, neologisms, and “word salad” observed in schizophrenia represent the active severing of the *FOXP2*-mediated linguistic tethers to the tribe.

When the metabolic cost of maintaining shared reality becomes too high, the Scout’s sensory apparatus outpaces its capacity for symbolic translation. The organism abandons shared language entirely to enter an unrestricted, low-latency “search mode” designed for environmental and thermodynamic survival. Because the modern environment fails to provide the high-density fuel necessary to exit this state, this adaptive search-mode is misidentified by modern diagnostics as permanent pathology.

Ultimately, the empirical persistence of this highly disruptive polygenic burden within the modern population fundamentally challenges the standard psychiatric “disease-deficit” framework. If the degradation of the *HCAR1/2* cooling circuit and lactate brake was simply a maladaptive biological error causing a catastrophic reduction in fitness, these alleles would have been subjected to rapid and heavy purifying selection. However, our constraint analysis demonstrates the exact opposite. The conservation of these alleles at ∼35% frequency indicates that the human population actively maintained these variants. We posit that this balancing selection preserves the high-risk, high-reward Vanguard Scout as a necessary evolutionary insurance policy—sacrificing baseline thermodynamic stability to maintain a reservoir of extreme associative plasticity during periods of severe environmental and metabolic volatility.

## Methods

### Data Sources and Cohort Acquisition

To evaluate the shared genomic architecture bridging the psychiatric spectrum and extreme cognitive plasticity, GWAS summary statistics were acquired from four primary cohorts. To operationalize the high-plasticity phenotype, data representing scientific occupational creativity (the “Creative Scientist” cohort) were obtained from the UK Biobank (GCST90444393) ^11^. For the severe psychiatric phenotypes, Schizophrenia (SCZ) summary statistics were obtained from the Psychiatric Genomics Consortium (PGC) Wave 3 dataset ^12^. Bipolar Disorder (BIP) summary statistics were acquired from the recent PGC 2024 meta-analysis ^14^. To assess structural variance across the broader neurodevelopmental spectrum, Autism Spectrum Disorder (ASD) data were sourced from the iPSYCH-PGC meta-analysis ^15^.

#### Targeted Locus Extraction and Genomic Coordinate Standardization

To isolate and quantify the polygenic burden across the posited “Vanguard Engine” network, we targeted ten prespecified candidate loci identified as critical high-voltage and thermodynamic hardware: *CACNA1C, CACNB2, GRIN2A, BDNF, FOXP2, PDHB, HCAR1/2, HDAC2, SNAP91*, and *DRD2*. The extraction of summary statistics for these specific gene bodies was executed using customized Unix-based Bash pipelines. To accurately capture local regulatory enhancers and structural variants without introducing excessive background noise, coordinate windows were dynamically tailored to each locus (spanning approximately 40 kb to 2 Mb) using standard high-throughput text-processing utilities. This generated the targeted locus datasets for the Creative Scientist **(Supplemental Data 2)**, Schizophrenia **(Supplemental Data 3)**, Bipolar Disorder **(Supplemental Data 4)**, and Autism Spectrum Disorder **(Supplemental Data 5)** cohorts.

Because the primary GWAS datasets were natively mapped to disparate genomic assemblies across their respective publication dates (e.g., GRCh37/hg19 versus GRCh38/hg38), cross-cohort spatial standardization was strictly enforced prior to comparative analysis. Rather than subjecting the summary statistics to absolute base-pair coordinate translation—which can introduce interpolation artifacts or variant dropout—structural variance across disparate assemblies was resolved via geometric normalization. Variant coordinates were mathematically mapped relative to their precise 5’ to 3’ position (expressed as a percentage) within the boundaries of their respective gene bodies. This spatial normalization established an assembly-agnostic framework, bypassing the need for raw coordinate conversion while ensuring absolute structural alignment during the generation of the cross-cohort comparative architectures and subsequent visual overlays.

#### Polygenic Burden and Locus-Zoom Analysis

To specifically evaluate the distal structural variance and regulatory burden within the cooling circuit, we performed an expanded, wide-regulatory search of the *HCAR1/2* locus. We extracted a wide 600 kb genomic window (representing an extended ±300 kb flanking region) centered on the *HCAR1/2* tandem loci (Chromosome 12: 122.8 Mb – 123.4 Mb; hg19 coordinates) from each primary GWAS dataset. Data extraction and formatting were performed using standard Unix text-processing utilities (awk, grep). Locus-zoom Manhattan plots were generated using Python with the matplotlib library, aligning the significance landscapes (−log_10_(*P*)) across the psychiatric spectrum. The *HCAR2* gene body (123.180 Mb – 123.190 Mb) was mapped as the structural anchor to contextualize the extended regulatory burden.

### Evolutionary Constraint and Balancing Selection

To establish an ancestral baseline for evolutionary constraint, minor allele frequencies (MAF) spanning the tandem *HCAR1/2* loci were retrieved from the 1000 Genomes Project (Phase 3; European Cohort) **(Supplemental Data 6)**. Data extraction was executed by streaming compressed variant call format (VCF) indices directly from the National Center for Biotechnology Information (NCBI) FTP mirror utilizing bcftools. The extracted ancestral frequencies were subsequently merged with the empirical Schizophrenia summary statistics (PGC Wave 3) using the pandas library in Python. To ensure absolute variant concordance, the datasets were cross-indexed strictly by their standardized genomic coordinates (hg19 base position). For the constraint analysis, the core structural variants (“Vanguard Hits”) were explicitly defined using a significance threshold of *P* < 1 × 10^-4^. The empirical cohort MAFs of these highly significant variants were then plotted against the ancestral 1000 Genomes baseline **(Fig. 3)**. The resulting distribution was evaluated relative to the neutral genetic drift diagonal (*y = x*) to mathematically distinguish between signatures of rapid purifying (negative) selection and sustained balancing selection within the human population.

### Archaic Divergence and Pan-Human Conservation Mapping

To subsequently investigate population-specific evolutionary trajectories across the broader network, minor allele frequencies for the core Vanguard structural variants were retrieved from the 1000 Genomes Project (Phase 3). Data extraction was executed via a custom programmatic pipeline querying the Ensembl REST API (https://rest.ensembl.org) **(Supplemental Data 7)**. This automated pipeline systematically interrogated the highly significant index variants (*P* < 1 × 10^-4^) across the ten core Vanguard loci, extracting specific allelic frequencies for the African (AFR), European (EUR), and East Asian (EAS) super-populations. To ensure absolute variant concordance, the script algorithmically resolved potential strand flips between the input GWAS summary statistics and the Ensembl reference genome.

To visualize evolutionary divergence and test for archaic introgression, the extracted ancestral frequencies were merged with the empirical Schizophrenia summary statistics (PGC Wave 3). The African (AFR) super-population minor allele frequencies were plotted against the European (EUR) super-population frequencies **(Fig. 4)**. The resulting distribution was evaluated relative to the neutral ancestral baseline diagonal (*y = x*). This spatial mapping allowed us to test for “Ice Age Introgression” variants (defined strictly as AFR frequency < 5% and EUR frequency > 20%). By mathematically ruling out these isolated post-migration selection pressures, this pipeline confirmed the deep, pan-human ancestral conservation of the Vanguard loci.

### Structural Mapping of the HCAR2 Receptor

To contextualize the kinetic and thermodynamic implications of the observed polygenic burden, a two-dimensional domain map of the *HCAR2* 7-transmembrane (7TM) G-protein-coupled receptor was constructed. This topological architecture was modeled upon the established high-resolution structural elucidation of the receptor ^17^. Specifically, the orthosteric binding pocket for BHB, niacin, and MMF—comprising transmembrane helices TM3, TM6, and TM7—and the intracellular *G_i_*-coupling domain at intracellular loop 3 (ICL3) were positionally isolated to visualize the functional thermodynamic axis. The massive regulatory polygenic burden identified in the severe psychiatric cohorts was conceptually superimposed onto this schematic as a structural fracture, visually representing the severe transcriptional downregulation and subsequent kinetic decay of the Vanguard lactate brake and cooling circuit.

## Acknowledgments

The author thanks stimulating discussions with Abraham Palmer at University of California, San Diego and Sean Crosson at Michigan State University. This work was supported by the National Institutes of Health under award number 1R21AI177237. The content is solely the responsibility of the author and does not necessarily represent the official views of the National Institutes of Health.

## Author Contributions

B.A.K. is the sole author of this manuscript. B.A.K. conceived the theoretical framework, executed the computational genomic analyses, performed the structural biophysical mapping, and wrote the manuscript.

## Data Availability

All primary data utilized in this study are derived from publicly available, de-identified genomic datasets. Schizophrenia, Bipolar Disorder, Autism Spectrum Disorder GWAS summary statistics are available via the Psychiatric Genomics Consortium (PGC) data portal (https://pgc.unc.edu). Scientific Occupational Creativity GWAS data are accessible via the UK Biobank repository (GCST90444393). Ancestral allele frequency data were retrieved from the 1000 Genomes Project Phase 3 data browser (https://www.internationalgenome.org).

## Code Availability

Custom Unix-based Bash pipelines and Python scripts utilized for targeted locus extraction, geometric coordinate normalization, and figure generation are publicly available via Zenodo (DOI: 10.5281/zenodo.20689820).

## Declaration of AI and AI-Assisted Technologies in the Writing Process

During the preparation of this work, the author used Google Gemini and Google Search to assist with literature research, refine manuscript text for clarity and readability, and optimize custom computational code. After using these tools/services, the author reviewed and edited the content as needed and takes full responsibility for the ultimate content and integrity of the publication.

