## Supplemental Figures and Table for "Metabolic Gating and the Evolution of Human Cognitive Plasticity: A Comparative Genomic Analysis"

Bryan A. Krantz

Department of Microbial Pathogenesis, School of Dentistry, University of Maryland, Baltimore, 650 W. Baltimore Street, Baltimore, MD 21201, U.S.A.

 (BAK)

**Running title:** Metabolic Evolution of Plasticity

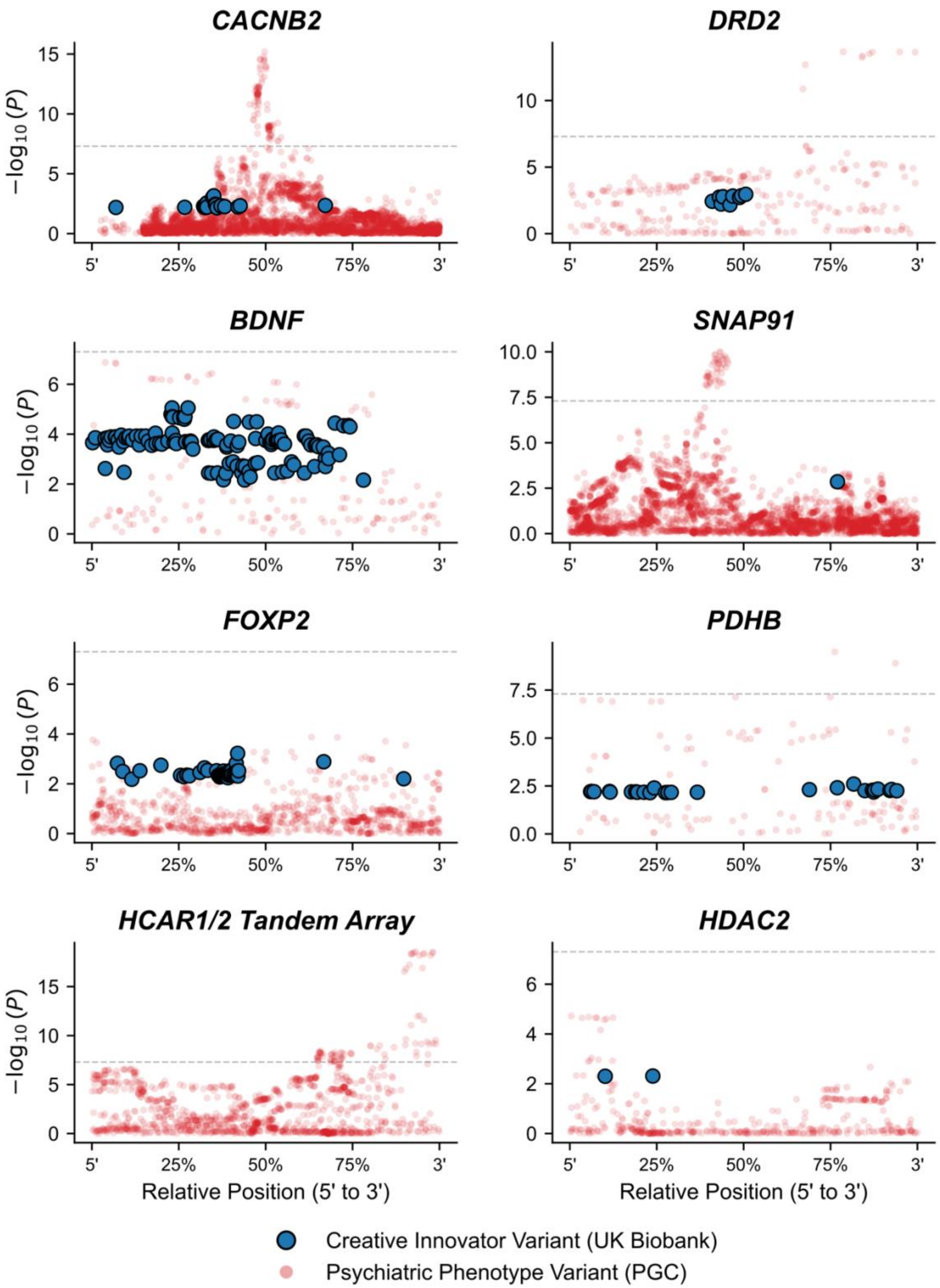

CACNB2
DRD2
BDNF
SNAP91
FOXP2
PDHB
HCAR1/2 Tandem Array
HDAC2
−log10(P)
5'
25%
50%
75%
3'
Relative Position (5' to 3')
Creative Innovator Variant (UK Biobank)
Psychiatric Phenotype Variant (PGC)

**Supplemental Figure S1: Extended Regional Genomic Burden of the Vanguard Engine and Metabolic Governors.** Functional locus plots detailing the geometric overlap of targeted structural variance across the remaining eight critical loci of the Vanguard architecture. The data compares the high-impact variants of the Scientific Creativity cohort (solid blue foreground; UK Biobank GCST90444393) against the dense polygenic burden of the Schizophrenia cohort (translucent red background; PGC Wave 3). To resolve spatial discrepancies between genome assemblies (hg19 vs. hg38), variant coordinates were geometrically normalized and plotted relative to their 5' to 3' position within the respective gene bodies. This extended inventory demonstrates shared structural vulnerability across auxiliary calcium modulation (*CACNB2*), neuromodulatory gating (*DRD2*), synaptic plasticity and vesicular transmission (*BDNF*, *SNAP91*), the cortico-striatal linguistic tether (*FOXP*2), and the complete metabolic and epigenetic governor circuit (*PDHB*, *HCAR1/2*, *HDAC2*). Grey dashed lines indicate the standard genome-wide significance threshold ($P < 5\times10^{-8}$). Note: Disparities in visual variant density reflect the inherent statistical power differences between the heavily powered psychiatric mega-GWAS and the targeted creativity cohort.

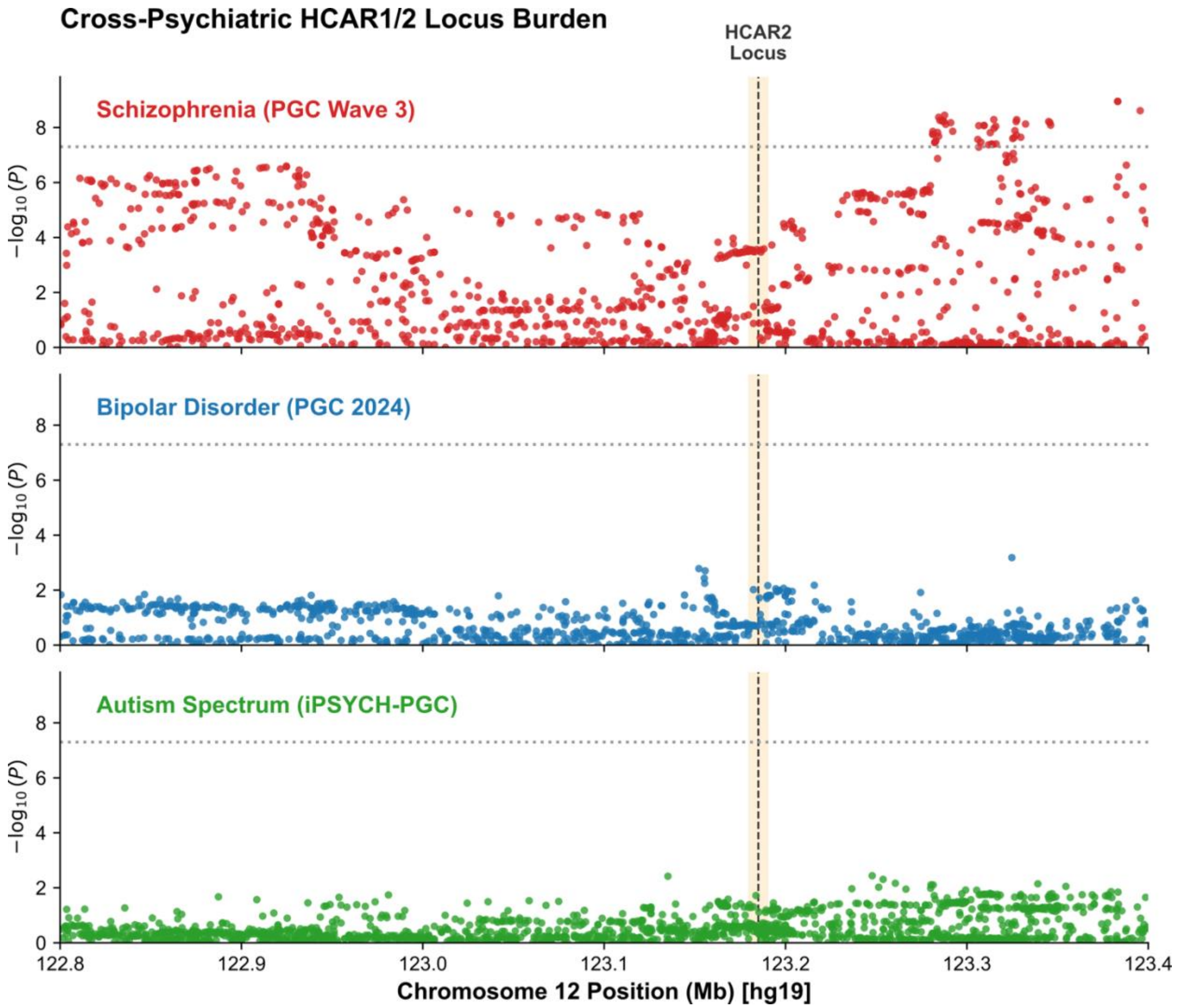

**Supplemental Figure S2: Conserved Polygenic Burden Across the Tandem HCAR1/2 Loci.** A comparative locus-zoom (Manhattan) analysis of the *HCAR1/2* wide regulatory and coding regions (Chromosome 12; hg19 coordinates) across three primary neurodevelopmental/psychiatric cohorts. The data demonstrate a conserved, severity-dependent structural burden within the cooling circuit. Schizophrenia (top panel, PGC Wave 3) exhibits a massive polygenic signal dominating the locus. Bipolar Disorder (middle panel, PGC 2024) presents a moderate, overlapping burden, while Autism Spectrum Disorder (bottom panel, iPSYCH) demonstrates minimal structural variance. The shaded region denotes the functional gene body of the *HCAR2* (GPR109A) receptor.

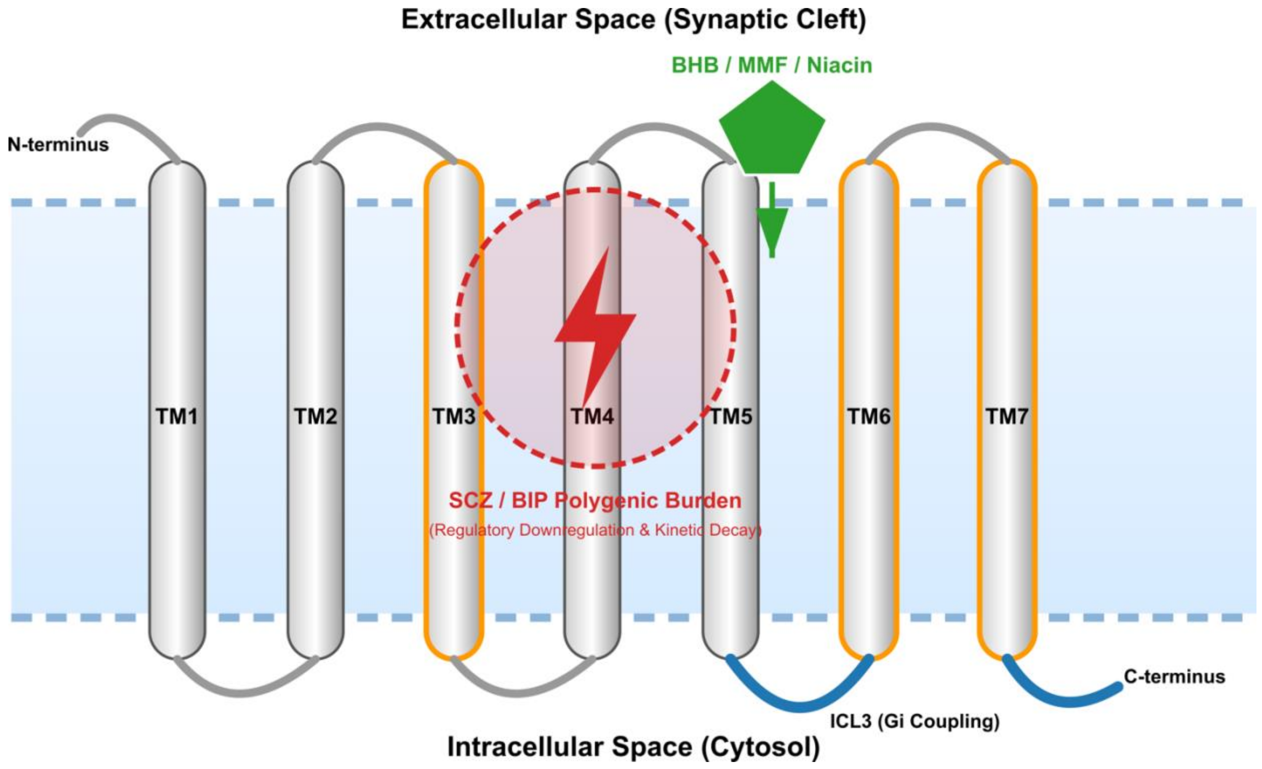

**Supplemental Figure S3: Biophysical and Functional Domain Map of the HCAR2 Cooling Circuit.** A 2D structural schematic of the 7-transmembrane (7TM) G-protein-coupled receptor HCAR2 (GPR109A). The schematic highlights the critical thermodynamic axis between the extracellular orthosteric binding pocket (green) and the intracellular $G_i$ coupling domain at Intracellular Loop 3 (ICL3, blue). The binding pocket serves as the primary sensor for ancestral ketogenic fuels (β-hydroxybutyrate), niacin, and potent neuro-inflammatory therapeutics (monomethyl fumarate (MMF)). The superimposed structural fracture overlay (red) represents the functional impact of the dense intronic and regulatory polygenic burden identified in the Schizophrenia (SCZ) and Bipolar Disorder (BIP) cohorts. Because these conserved variants drive transcriptional downregulation and kinetic decay rather than localized amino acid substitutions, the resulting pathology is a systemic kinetic uncoupling: the receptor fails to translate the presence of the ligand into the necessary $G_i$ -mediated neuro-inflammatory quenching cascade, precipitating the thermal overload of the Vanguard Engine.

**Supplementary Table S1: Lead Regulatory Variants and Polygenic Burden at the *HCAR1/2* Locus.**

| Disorder | rsID | Chromosome | Position (hg19) | Effect Allele | Other Allele | *P*-Value | Beta | SE |
|---|---|---|---|---|---|---|---|---|
| SCZ | rs80024005 | 12 | 123383147 | G | T | 1.12E-09 | -0.0952 | 0.0156 |
| SCZ | rs76446716 | 12 | 123383148 | C | T | 1.12E-09 | -0.0952 | 0.0156 |
| SCZ | rs75438052 | 12 | 123395555 | A | C | 2.44E-09 | -0.0839 | 0.0141 |
| SCZ | rs58182048 | 12 | 123287708 | C | T | 3.56E-09 | -0.0861 | 0.0146 |
| SCZ | rs75482067 | 12 | 123284472 | G | A | 4.22E-09 | -0.0853 | 0.0145 |
| SCZ | rs80029885 | 12 | 123314853 | G | A | 5.14E-09 | -0.0853 | 0.0146 |
| SCZ | rs12298664 | 12 | 123327445 | T | C | 5.16E-09 | -0.0847 | 0.0145 |
| SCZ | rs12321131 | 12 | 123285124 | G | A | 5.51E-09 | -0.0847 | 0.0145 |
| SCZ | rs79775390 | 12 | 123286774 | C | T | 5.58E-09 | -0.0847 | 0.0145 |
| SCZ | rs11060065 | 12 | 123286491 | A | G | 5.59E-09 | -0.0847 | 0.0145 |
| **Disorder** | **rsID** | **Chromosome** | **Position (hg19)** | **Effect Allele** | **Other Allele** | ***P*-Value** | **Odds Ratio (OR)** | **SE** |
| BIP | rs117703564 | 12 | 123324810 | A | G | 6.57E-04 | 1.144 | 0.0395 |
| BIP | rs4759370 | 12 | 123152216 | G | A | 1.65E-03 | 0.9539 | 0.015 |
| BIP | rs34371863 | 12 | 123155683 | C | T | 1.98E-03 | 0.9563 | 0.0144 |
| BIP | rs35576166 | 12 | 123155218 | T | A | 3.69E-03 | 0.9591 | 0.0144 |
| **Disorder** | **rsID** | **Chromosome** | **Position (hg19)** | **Effect Allele** | **Other Allele** | ***P*-Value** | **Odds Ratio (OR)** | **SE** |
| ASD | rs11384734 | 12 | 123247791 | GA | G | 3.64E-03 | 0.9458 | 0.0192 |
| ASD | rs138608943 | 12 | 123135148 | A | G | 3.80E-03 | 1.1877 | 0.0594 |
| ASD | rs34056825 | 12 | 123253798 | T | TAC | 4.92E-03 | 1.0724 | 0.0249 |

This summary table details the top 10 most significant structural and regulatory variants localized to the *HCAR1/2* braking and cooling circuits (Chromosome 12; hg19 coordinates) across the three primary psychiatric and neurodevelopmental cohorts. Variants are categorized by their respective clinical phenotype: Schizophrenia (SCZ; PGC Wave 3), Bipolar Disorder (BIP; PGC 2024), and Autism Spectrum Disorder (ASD; iPSYCH). Summary statistics include the designated disorder, rsID, absolute genomic position, effect/reference alleles, association *P*-values, and effect sizes (Beta or Odds Ratio) with standard errors (SE). The exhaustive compilation of all nominally significant structural variants ($P < 5 \times 10^{-3}$) across these cohorts—detailing the massive structural divergence in SCZ, the intermediate burden in BIP, and the minimal variance in ASD—is provided as a separate, downloadable Excel workbook (**Supplemental Data 1**).
