## Supplemental Data 2 for "Metabolic Gating and the Evolution of Human Cognitive Plasticity: A Comparative Genomic Analysis": bash-Creativity-GCST90444393.sh

```
# DRD2 (hg38: chr11)
awk 'NR==1 {print} $1=="11" && $2>=113350000 && $2<=113550000 {print}' GCST90444393.tsv > Creativity_DRD2_hits.tsv

# FOXP2 (hg38: chr7)
awk 'NR==1 {print} $1=="7" && $2>=114000000 && $2<=114800000 {print}' GCST90444393.tsv > Creativity_FOXP2_hits.tsv

# BDNF (hg38: chr11)
awk 'NR==1 {print} $1=="11" && $2>=27600000 && $2<=27800000 {print}' GCST90444393.tsv > Creativity_BDNF_hits.tsv

# HCAR2 (hg38: chr12)
awk 'NR==1 {print} $1=="12" && $2>=122600000 && $2<=122800000 {print}' GCST90444393.tsv > Creativity_HCAR2_hits.tsv

# PDHB (hg38: chr3)
awk 'NR==1 {print} $1=="3" && $2>=53350000 && $2<=53550000 {print}' GCST90444393.tsv > Creativity_PDHB_hits.tsv

# CACNA1C (hg38: chr12)
awk 'NR==1 {print} $1=="12" && $2>=1950000 && $2<=2550000 {print}' GCST90444393.tsv > Creativity_CACNA1C_hits.tsv

# CACNB2 (hg38: chr10)
awk 'NR==1 {print} $1=="10" && $2>=18150000 && $2<=18750000 {print}' GCST90444393.tsv > Creativity_CACNB2_hits.tsv

# GRIN2A (hg38: chr16)
awk 'NR==1 {print} $1=="16" && $2>=9650000 && $2<=10250000 {print}' GCST90444393.tsv > Creativity_GRIN2A_hits.tsv

# SNAP91 (hg38: chr6 - included here for your complete record)
awk 'NR==1 {print} $1=="6" && $2>=83350000 && $2<=83650000 {print}' GCST90444393.tsv > Creativity_SNAP91_hits.tsv

# HDAC2 (hg38: chr6)
awk 'NR==1 {print} $1=="6" && $2>=114200000 && $2<=114400000 {print}' GCST90444393.tsv > Creativity_HDAC2_hits.tsv
```
