## Supplemental Data 2 for "Metabolic Gating and the Evolution of Human Cognitive Plasticity: A Comparative Genomic Analysis": README.docx

**README: Supplementary Data 2**

**Dataset:** Full .tsv.zip composite file of gene series queries for the Creative Scientists cohort. **Associated Manuscript:** Metabolic Gating and the Evolution of Human Cognitive Plasticity: A Comparative Genomic Analysis.

**1. Dataset Overview**

This archive contains the targeted genome-wide association study (GWAS) summary statistics for the 10 prespecified candidate loci comprising the "Vanguard Engine" network (*CACNA1C, CACNB2, GRIN2A, BDNF, FOXP2, PDHB, HCAR2, HDAC2, SNAP91*, and *DRD2*). The data represents the "Scientific Occupational Creativity" cohort, serving as the high-plasticity, homeostatic phenotype in the comparative analysis.

**2. Parent Genomic Data Origin**

The raw, genome-wide summary statistics were acquired from the **UK Biobank (UKB)**.

- **Phenotype:** Scientific Occupational Creativity
- **Reference Study:** Li et al., 2024 (*Commun Biol*)
- **NHGRI-EBI GWAS Catalog Accession:** GCST90444393
- **Primary Repository:** Data is publicly hosted and retrievable via the GWAS Catalog FTP server.

**3. Genome Assembly Architecture & Spatial Normalization**

- **Native Assembly:** The parent summary statistics for GCST90444393 natively map to the **GRCh38 (hg38)** human reference genome assembly.
- **Standardization:** Because the comparative psychiatric datasets (PGC/iPSYCH) map to GRCh37 (hg19), structural variance comparisons across disparate assemblies were resolved via geometric normalization. Variant coordinates were mapped relative to their precise 5' to 3' position (percentage) within the respective gene bodies, bypassing the need for raw coordinate translation while preserving absolute spatial accuracy.

**4. Extraction Methodology**

Data extraction was executed utilizing a customized Unix-based Bash pipeline (bash-Creativity-GCST90444393.sh). Standard high-throughput text-processing utilities (awk, grep) were deployed to stream the parent GWAS dataset and isolate the specific gene bodies of the targeted loci.

**Extraction Parameters:**

- Each locus query includes a robust flanking window of **±300 kb** from the established 5' and 3' transcription start/end sites.
- This windowing strategy was mathematically enforced to comprehensively capture distal regulatory enhancers, promoter regions, and structural variants influencing transcriptional control.

**5. File Structure**

The unzipped .tsv composite file contains the following harmonized columns derived from the parent GWAS Catalog summary statistics:

- CHR: Chromosome (hg19)
- POS: Base pair position (hg19)
- SNP: Variant identifier (rsID)
- Effect_Allele / Other_Allele: Evaluated alleles
- Beta: Effect size
- SE: Standard Error
- P_Value: Association significance
- Gene_Target: The specific Vanguard locus assigned to the extracted variant
