## Supplemental Data 3 for "Metabolic Gating and the Evolution of Human Cognitive Plasticity: A Comparative Genomic Analysis": bash-collection-SCZ-PGC.sh

```
zgrep -v "^##" PGC3_SCZ_wave3.primary.autosome.public.v3.vcf.tsv.gz | awk 'NR==1 {print} $1=="5" && $3>=73800000 && $3<=73860000 {print}' > PRKAA1_hits.tsv
zgrep -v "^##" PGC3_SCZ_wave3.primary.autosome.public.v3.vcf.tsv.gz | awk 'NR==1 {print} $1=="1" && $3>=56900000 && $3<=56980000 {print}' > PRKAA2_hits.tsv
zgrep -v "^##" PGC3_SCZ_wave3.primary.autosome.public.v3.vcf.tsv.gz | awk 'NR==1 {print} $1=="11" && $3>=4000000 && $3<=6000000 {print}' > Olfactory_hits.tsv
zgrep -v "^##" PGC3_SCZ_wave3.primary.autosome.public.v3.vcf.tsv.gz | awk 'NR==1 {print} $1=="12" && $3>=10800000 && $3<=11500000 {print}' > Bitter_Toxin_hits.tsv
zgrep -v "^##" PGC3_SCZ_wave3.primary.autosome.public.v3.vcf.tsv.gz | awk 'NR==1 {print} $1=="1" && $3>=16000000 && $3<=16500000 {print}' > TAS1R1_Umami_hits.tsv
zgrep -v "^##" PGC3_SCZ_wave3.primary.autosome.public.v3.vcf.tsv.gz | awk 'NR==1 {print} $1=="1" && $3>=1200000 && $3<=1350000 {print}' > TAS1R3_Umami_hits.tsv
zgrep -v "^##" PGC3_SCZ_wave3.primary.autosome.public.v3.vcf.tsv.gz | awk 'NR==1 {print} $1=="12" && $3>=53290000 && $3<=53310000 {print}' > Null_Keratin_hits.tsv
zgrep -v "^##" PGC3_SCZ_wave3.primary.autosome.public.v3.vcf.tsv.gz | awk 'NR==1 {print} $1=="12" && $3>=59980000 && $3<=60180000 {print}' > BHB_MCT2_hits.tsv
zgrep -v "^##" PGC3_SCZ_wave3.primary.autosome.public.v3.vcf.tsv.gz | awk 'NR==1 {print} $1=="12" && $3>=123150000 && $3<=123200000 {print}' > BHB_HCAR2_hits.tsv
zgrep -v "^##" PGC3_SCZ_wave3.primary.autosome.public.v3.vcf.tsv.gz | awk 'NR==1 {print} $1=="5" && $3>=141000000 && $3<=141020000 {print}' > BHB_HDAC3_hits.tsv
zgrep -v "^##" PGC3_SCZ_wave3.primary.autosome.public.v3.vcf.tsv.gz | awk 'NR==1 {print} $1=="19" && $3>=35840000 && $3<=35860000 {print}' > BHB_FFAR3_hits.tsv
zgrep -v "^##" PGC3_SCZ_wave3.primary.autosome.public.v3.vcf.tsv.gz | awk 'NR==1 {print} $1=="11" && $3>=113250000 && $3<=113350000 {print}' > Symbol_DRD2_hits.tsv
zgrep -v "^##" PGC3_SCZ_wave3.primary.autosome.public.v3.vcf.tsv.gz | awk 'NR==1 {print} $1=="7" && $3>=113700000 && $3<=114350000 {print}' > Symbol_FOXP2_hits.tsv
zgrep -v "^##" PGC3_SCZ_wave3.primary.autosome.public.v3.vcf.tsv.gz | awk 'NR==1 {print} $1=="11" && $3>=27650000 && $3<=27750000 {print}' > Symbol_BDNF_hits.tsv
zgrep -v "^##" PGC3_SCZ_wave3.primary.autosome.public.v3.vcf.tsv.gz | awk 'NR==1 {print} $1=="7" && $3>=137700000 && $3<=137740000 {print}' > Metabolism_PDK4_Lock_hits.tsv
zgrep -v "^##" PGC3_SCZ_wave3.primary.autosome.public.v3.vcf.tsv.gz | awk 'NR==1 {print} $1=="3" && $3>=53390000 && $3<=53460000 {print}' > Metabolism_PDHB_Door_hits.tsv
zgrep -v "^##" PGC3_SCZ_wave3.primary.autosome.public.v3.vcf.tsv.gz | awk 'NR==1 {print} $1=="16" && $3>=9500000 && $3<=10500000 {print}' > Symbol_GRIN2A_hits.tsv
zgrep -v "^##" PGC3_SCZ_wave3.primary.autosome.public.v3.vcf.tsv.gz | awk 'NR==1 {print} $1=="12" && $3>=1800000 && $3<=3000000 {print}' > Symbol_CACNA1C_hits.tsv
zgrep -v "^##" PGC3_SCZ_wave3.primary.autosome.public.v3.vcf.tsv.gz | awk 'NR==1 {print} $1=="10" && $3>=18000000 && $3<=19500000 {print}' > Symbol_CACNB2_hits.tsv
zgrep -v "^##" PGC3_SCZ_wave3.primary.autosome.public.v3.vcf.tsv.gz | awk 'NR==1 {print} $1=="6" && $3>=83500000 && $3<=85500000 {print}' > Symbol_SNAP91_hits.tsv
zgrep -v "^##" PGC3_SCZ_wave3.primary.autosome.public.v3.vcf.tsv.gz | awk 'NR==1 {print} $1=="12" && $3>=6630000 && $3<=6660000 {print}' > Null_GAPDH_hits.tsv
zgrep -v "^##" PGC3_SCZ_wave3.primary.autosome.public.v3.vcf.tsv.gz | awk 'NR==1 {print} $1=="6" && $3>=114200000 && $3<=114350000 {print}' > BHB_HDAC2_hits.tsv
zgrep -v "^##" PGC3_SCZ_wave3.primary.autosome.public.v3.vcf.tsv.gz | awk 'NR==1 {print} $1=="12" && $3>=122800000 && $3<=123400000 {print}' > BHB_HCAR2_wide_regulatory.tsv
```
