## Supplemental Data 3 for "Metabolic Gating and the Evolution of Human Cognitive Plasticity: A Comparative Genomic Analysis": README.docx

**README: Supplementary Data 3**

**Dataset:** Full .tsv.zip composite file of gene series queries for Schizophrenia (PGC). **Associated Manuscript:** Metabolic Gating and the Evolution of Human Cognitive Plasticity: A Comparative Genomic Analysis.

**1. Dataset Overview**

This archive contains the targeted genome-wide association study (GWAS) summary statistics mapping structural variance in the severe psychiatric phenotype (Schizophrenia). It includes queries for the 10 prespecified candidate loci comprising the "Vanguard Engine" network, expanded exploratory loci for sensory and metabolic gating, and standard null control loci to establish baseline genomic variance.

**2. Parent Genomic Data Origin**

The raw, genome-wide summary statistics were acquired from the **Psychiatric Genomics Consortium (PGC)**.

- **Cohort:** Schizophrenia (SCZ) Wave 3
- **Reference Study:** Trubetskoy et al., 2022 (*Nature*)
- **Primary Repository:** Data is publicly hosted and retrievable via the PGC Data Portal.

**3. Genome Assembly Architecture & Coordinate Standardization**

- **Native Assembly:** The parent summary statistics for the PGC Wave 3 dataset natively map to the **GRCh37 (hg19)** human reference genome assembly.
- **Standardization:** All genomic coordinates (Chromosome and Base Pair Position) within this extracted .tsv dataset have been strictly verified against the **GRCh37/hg19** architecture to prevent assembly shifts and ensure precise variant-to-variant alignment across cross-cohort plotting.

**4. Extraction Methodology**

Data extraction was executed utilizing a customized Unix-based Bash pipeline (bash-collection-SCZ-PGC.sh). Standard text-processing utilities were deployed to stream the parent GWAS dataset and isolate the specific genomic coordinates of the targeted loci.

**Standard Extraction Parameters:**

- The core locus queries include a robust flanking window of **±300 kb** from the established 5' and 3' transcription start/end sites to comprehensively capture distal regulatory enhancers and structural variants influencing transcriptional control.

**Exceptions & Wide Regulatory Mapping:**

- BHB_HCAR2_wide_regulatory.tsv: This file represents an independent, expanded search window specifically deployed to capture deep regulatory sequence space beyond the standard boundary constraints for the *HCAR2* (GPR109A) locus.

**5. File Manifest & Structure**

The unzipped .tsv composite file contains the following extracted datasets, categorized by their functional testing parameters:

A. The Core Vanguard Hardware & Governors

- Symbol_CACNA1C_hits.tsv
- Symbol_CACNB2_hits.tsv
- Symbol_GRIN2A_hits.tsv
- Symbol_DRD2_hits.tsv
- Symbol_SNAP91_hits.tsv
- Symbol_BDNF_hits.tsv
- Symbol_FOXP2_hits.tsv
- Metabolism_PDHB_Door_hits.tsv
- BHB_HCAR2_hits.tsv
- BHB_HDAC2_hits.tsv

**B. Extended Metabolic & Epigenetic Probes**

- BHB_HCAR2_wide_regulatory.tsv *(Expanded regulatory architecture)*
- BHB_HDAC3_hits.tsv
- BHB_FFAR3_hits.tsv
- BHB_MCT2_hits.tsv
- Metabolism_PDK4_Lock_hits.tsv
- PRKAA1_hits.tsv
- PRKAA2_hits.tsv

**C. Sensory & Olfactory Pivot Probes**

*(Investigating the Vanguard Scout's fuel detection and environmental anomaly circuits)*

- Olfactory_hits.tsv
- Bitter_Toxin_hits.tsv
- TAS1R1_Umami_hits.tsv
- TAS1R3_Umami_hits.tsv

**D. Standardized Null Controls** *(Housekeeping and structural baseline loci)*

- Null_GAPDH_hits.tsv
- Null_Keratin_hits.tsv

**6. Column Formatting**

All .tsv files maintain standardized column headers derived from the PGC summary statistics:

- CHR: Chromosome (hg19)
- POS: Base pair position (hg19)
- SNP: Variant identifier (rsID)
- A1 / A2: Effect Allele / Reference Allele
- BETA / OR: Effect size / Odds Ratio
- SE: Standard Error
- P: Association significance
