## Supplemental Data 4 for "Metabolic Gating and the Evolution of Human Cognitive Plasticity: A Comparative Genomic Analysis": bipolar_HCAR2_bash.sh

```
# Narrow focused search
gzcat bip2024_eur_no23andMe.gz | awk 'NR==1 {print} $2=="12" && $3>=123150000 && $3<=123200000 {print}' > BIP_HCAR2_hits.tsv

# Wide redulatory search
gzcat bip2024_eur_no23andMe.gz | awk 'NR==1 {print} $2=="12" && $3>=122800000 && $3<=123400000 {print}' > BIP_HCAR2_wide_regulatory.txt
```
