## Supplemental Data 4 for "Metabolic Gating and the Evolution of Human Cognitive Plasticity: A Comparative Genomic Analysis": README.docx

**README: Supplementary Data 4**

**Dataset:** Full .zip archive of targeted *HCAR2* gene queries for Bipolar Disorder (PGC). **Associated Manuscript:** Metabolic Gating and the Evolution of Human Cognitive Plasticity: A Comparative Genomic Analysis.

**1. Dataset Overview**

This archive contains the targeted genome-wide association study (GWAS) summary statistics mapping structural variance across the *HCAR2* locus in the Bipolar Disorder (BIP) phenotype.

**Format Note:** Unlike the tab-separated (.tsv) datasets provided for the Schizophrenia and Creativity cohorts, the parent summary statistics for this dataset were provided in the Ricopili daner format. Consequently, the extracted files herein are strictly **space-delimited** and have been designated with a .txt extension to accurately reflect this structural formatting and prevent parsing errors.

**2. Parent Genomic Data Origin**

The raw, genome-wide summary statistics were acquired from the **Psychiatric Genomics Consortium (PGC)**.

- **Cohort:** Bipolar Disorder Meta-analysis
- **Reference Study:** O'Connell et al.
- **Data Steward:** Kevin O'Connell
- **Primary Repository:** Data is publicly hosted and retrievable via the PGC Data Portal.

**3. Genome Assembly Architecture & Source Quality Control**

- **Native Assembly:** The parent summary statistics natively map to the **GRCh37 (hg19)** human reference genome assembly.
- **Quality Control (QC) Parameters:** The parent data underwent rigorous pre-extraction filtering by the PGC. The following QC parameters are inherent to the data contained in this archive:
  - Variants present in < 75% of the total effective sample size were removed.
  - Variants were retained strictly if the cohort minor allele frequency (MAF) > 1% and the minor allele count (MAC) > 10 in either cases or controls.
  - The DENTIST algorithmic tool was employed by the source authors to detect and exclude problematic variants prior to release.
  - Allele frequencies provided correspond to A1 from the HRC reference panel corresponding to the meta-analysis ancestry.

**4. Extraction Methodology**

Data extraction was executed utilizing a customized Unix-based Bash pipeline (bipolar_HCAR2_bash.sh). Standard text-processing utilities were deployed to stream the space-delimited parent GWAS dataset and isolate the specific genomic coordinates of the *HCAR2* locus.

**Extraction Parameters:**

- **Standard Window:** The primary query utilized a robust flanking window of **±300 kb** from the established transcription start/end sites to capture standard promoter elements and structural variants.
- **Wide Regulatory Mapping:** An independent, expanded search window was subsequently deployed specifically to capture deep regulatory sequence space beyond the standard boundary constraints for the *HCAR2* (GPR109A) locus.

**5. File Manifest & Structure**

The unzipped archive contains the following extracted space-delimited (.txt) datasets:

- BIP_HCAR2_hits.txt: The standard structural/regulatory footprint for the *HCAR2* locus.
- BIP_HCAR2_wide_regulatory.txt: The expanded regulatory architecture mapping deep distal enhancers.

**Column Headers (Ricopili daner standard):** Includes standard mapping data (CHR, BP, SNP), allele tracking (A1, A2), and association metrics (OR, SE, P), among other standard PGC output metrics.
