## Supplemental Data 5 for "Metabolic Gating and the Evolution of Human Cognitive Plasticity: A Comparative Genomic Analysis": ASD_HCAR2_bash.sh

```
# Narrow focused HCAR2 search
gzcat iPSYCH-PGC_ASD_Nov2017.gz | awk 'NR==1 {print} $1=="12" && $3>=123150000 && $3<=123200000 {print}' > ASD_HCAR2_hits.tsv

# Wide regulatory HCAR2 search
gzcat iPSYCH-PGC_ASD_Nov2017.gz | awk 'NR==1 {print} $1=="12" && $3>=122800000 && $3<=123400000 {print}' > ASD_HCAR2_wide_regulatory.tsv
```
