## Supplemental Data 5 for "Metabolic Gating and the Evolution of Human Cognitive Plasticity: A Comparative Genomic Analysis": README.docx

**README: Supplementary Data 5**

**Dataset:** Full .tsv.zip archive of targeted *HCAR2* gene queries for Autism Spectrum Disorder (ASD). **Associated Manuscript:** Metabolic Gating and the Evolution of Human Cognitive Plasticity: A Comparative Genomic Analysis.

**1. Dataset Overview**

This archive contains the targeted genome-wide association study (GWAS) summary statistics mapping structural variance across the *HCAR2* locus in the Autism Spectrum Disorder (ASD) phenotype. This dataset serves as the neurodevelopmental baseline in the comparative cross-cohort structural analysis.

**2. Parent Genomic Data Origin**

The raw, genome-wide summary statistics were acquired from the **Psychiatric Genomics Consortium (PGC) and the iPSYCH Project**.

- **Cohort:** iPSYCH-PGC ASD Meta-analysis (Nov 2017 Release)
- **Reference Study:** Grove et al., 2019 (*Nature Genetics*)
- **Primary Repository:** Data is publicly hosted and retrievable via the PGC Data Portal.

**3. Genome Assembly Architecture**

- **Native Assembly:** The parent summary statistics for the iPSYCH-PGC 2017 dataset natively map to the **GRCh37 (hg19)** human reference genome assembly.
- **Standardization:** All genomic coordinates (Chromosome and Base Pair Position) within this extracted .tsv dataset have been strictly verified against the **GRCh37/hg19** architecture to ensure precise spatial alignment and variant concordance when plotted against the Schizophrenia and Bipolar cohorts.

**4. Extraction Methodology**

Data extraction was executed utilizing a customized Unix-based Bash pipeline (ASD_HCAR2_bash.sh). Standard high-throughput text-processing utilities were deployed to stream the parent GWAS dataset and isolate the specific genomic coordinates of the *HCAR2* locus.

**Extraction Parameters:**

- **Standard Window:** The primary query utilized a robust flanking window of **±300 kb** from the established *HCAR2* (GPR109A) transcription start/end sites to capture standard promoter elements and structural variants.
- **Wide Regulatory Mapping:** An independent, expanded search window was deployed to map the deep distal enhancer architecture and wide regulatory sequence space. This was methodologically required to ensure absolute comparative parity with the expanded regulatory queries utilized in the Schizophrenia and Bipolar Disorder analyses.

**5. File Manifest & Structure**

The unzipped archive contains the following extracted tab-separated (.tsv) datasets:

- ASD_HCAR2_hits.tsv: The standard structural and proximal regulatory footprint for the *HCAR2* locus.
- ASD_HCAR2_wide_regulatory.tsv: The expanded regulatory architecture capturing deep spatial boundaries.

**Column Formatting:** All .tsv files maintain standardized column headers derived from the PGC/iPSYCH summary statistics (e.g., CHR, SNP, BP, A1, A2, INFO, OR, SE, P).
