## Supplemental Data 6 for "Metabolic Gating and the Evolution of Human Cognitive Plasticity: A Comparative Genomic Analysis": README.docx

**README: Supplementary Data 6**

**Dataset:** Full .tsv.zip archive of ancestral allele frequencies for the *HCAR2* extended locus. **Associated Manuscript:** Metabolic Gating and the Evolution of Human Cognitive Plasticity: A Comparative Genomic Analysis.

**1. Dataset Overview**

This archive contains the baseline ancestral allele frequencies for the extended *HCAR2* regulatory window (Chromosome 12: 122.8 Mb – 123.4 Mb). This data was utilized to mathematically map evolutionary constraint, calculate purifying versus balancing selection signatures, and establish the population frequency baselines (Figure S4) for the structural variants identified in the psychiatric cohorts.

**2. Parent Genomic Data Origin**

The raw genotypic data was acquired directly from the **1000 Genomes Project**.

- **Cohort:** Phase 3 (Release 20130502)
- **Dataset:** ALL.chr12.phase3_shapeit2_mvncall_integrated_v5a.20130502.genotypes.vcf.gz
- **Primary Repository:** National Center for Biotechnology Information (NCBI) FTP Trace Archive.

**3. Genome Assembly Architecture**

- **Native Assembly:** The 1000 Genomes Phase 3 data maps natively to the **GRCh37 (hg19)** human reference genome assembly.
- **Concordance:** This assembly is perfectly synchronized with the hg19 coordinates utilized in the UK Biobank (Creativity) and PGC (Schizophrenia, Bipolar, ASD) cohort extractions, ensuring absolute variant-to-variant spatial alignment during the cross-cohort constraint analysis.

**4. Extraction Methodology**

Data extraction was executed utilizing bcftools to remotely query and stream the compressed Variant Call Format (VCF) index directly from the NCBI FTP mirror, bypassing the need for full-chromosome local downloads.

**Extraction Parameters & Command line:** The extraction targeted a precise 600 kb genomic window encapsulating the *HCAR2* gene body and its wide regulatory boundaries.

Bash

bcftools query -r 12:122800000-123400000 \

-f '%CHROM\t%POS\t%ID\t%REF\t%ALT\t%INFO/AF\t%INFO/EUR_AF\n' \

https://ftp-trace.ncbi.nih.gov/1000genomes/ftp/release/20130502/ALL.chr12.phase3_shapeit2_mvncall_integrated_v5a.20130502.genotypes.vcf.gz \

> 1000G_HCAR2_freqs.tsv

**5. File Manifest & Structure**

The unzipped archive contains a single tab-separated (.tsv) dataset:

- 1000G_HCAR2_freqs.tsv

**Column Formatting:** The columns directly correspond to the extraction flags utilized in the query, providing the necessary metrics for constraint mapping against the psychiatric cohorts:

- CHROM: Chromosome (hg19)
- POS: Base pair position (hg19)
- ID: Variant identifier (rsID)
- REF: Ancestral Reference Allele
- ALT: Alternate Allele
- AF: Global Allele Frequency (All Ancestries)
- EUR_AF: European Ancestry Allele Frequency
