## Supplemental Data 7 for "Metabolic Gating and the Evolution of Human Cognitive Plasticity: A Comparative Genomic Analysis": archaic_divergence_plotter.py

```
import pandas as pd
import matplotlib.pyplot as plt
import seaborn as sns
import matplotlib as mpl
import os

# ==========================================
# 0. Global Settings for Publication SVG Export
# ==========================================
mpl.rcParams['svg.fonttype'] = 'none'
plt.rcParams['font.family'] = 'sans-serif'
plt.rcParams['font.sans-serif'] = ['Arial']
plt.rcParams['font.size'] = 11

def generate_archaic_plot():
    print("--- Generating Publication-Quality Archaic Divergence Plot ---")
    
    input_file = "Split_HCAR_Archaic_Divergence.tsv"
    
    if not os.path.exists(input_file):
        print(f"⚠️ Error: Could not find '{input_file}'.")
        return
        
    try:
        df = pd.read_csv(input_file, sep='\t')
        print(f"Loaded {len(df)} Vanguard variants for plotting.")
    except Exception as e:
        print(f"Error loading file: {e}")
        return

    sns.set_theme(style="whitegrid", context="paper")
    
    # Widen the figure significantly to leave room for the massive legend
    fig, ax = plt.subplots(figsize=(10, 8)) 
    
    # Draw the Neutral Drift diagonal line
    ax.plot([-0.05, 1.05], [-0.05, 1.05], 'k--', alpha=0.5, zorder=1, label="Neutral Ancestral Baseline (Pan-Human)")
    
    # Highlight the "Smoking Gun" Archaic Introgression Zone (AFR < 5%, EUR > 20%)
    import matplotlib.patches as patches
    introgression_box = patches.Rectangle(
        (-0.05, 0.20), 0.10, 0.85, # (x, y), width, height
        linewidth=0, edgecolor='none', facecolor='red', alpha=0.08, zorder=0,
        label="Ice Age Introgression Zone"
    )
    ax.add_patch(introgression_box)

    scatter = sns.scatterplot(
        data=df, 
        x='AFR_Freq', 
        y='EUR_Freq', 
        hue='Locus', 
        palette='tab10', # tab10 perfectly accommodates up to 10 distinct loci
        s=90,            # Bubble size
        edgecolor='black',
        linewidth=0.5,
        alpha=0.85,
        zorder=3,
        ax=ax
    )
    
    ax.set_title("Pan-Human Ancestral Conservation of the Vanguard Cognitive Engine", 
                 fontsize=14, fontweight='bold', pad=15)
    ax.set_xlabel("African Allele Frequency (Ancestral Baseline)", 
                  fontsize=12, fontweight='bold')
    ax.set_ylabel("European Allele Frequency (Post-Migration Introgression)", 
                  fontsize=12, fontweight='bold')
    
    # Standardize axes from 0 to 1 (100% frequency) to show the full evolutionary spectrum
    ax.set_xlim(-0.05, 1.05)
    ax.set_ylim(-0.05, 1.05) 
    
    handles, labels = ax.get_legend_handles_labels()
    
    # Rebuild the legend to drop duplicate labels if seaborn added its own title
    legend = ax.legend(handles, labels, bbox_to_anchor=(1.05, 1), loc='upper left', 
                       frameon=True, title="Vanguard Network Loci", borderaxespad=0.)
    legend.get_title().set_fontweight('bold')

    # Ensure the layout doesn't chop off the external legend during export
    plt.tight_layout()
    
    svg_out = "Figure_Hypothesis_Archaic_Divergence.svg"
    png_out = "Figure_Hypothesis_Archaic_Divergence.png"
    
    plt.savefig(svg_out, format="svg", bbox_inches="tight")
    plt.savefig(png_out, format="png", dpi=300, bbox_inches="tight")
    
    print("✅ Success! Figure generated.")
    print(f"📁 Vector SVG: {svg_out}")
    print(f"📁 High-Res PNG: {png_out}")

if __name__ == "__main__":
    generate_archaic_plot()
```
