## Supplemental Data 7 for "Metabolic Gating and the Evolution of Human Cognitive Plasticity: A Comparative Genomic Analysis": README.txt

# Supplemental Dataset 7: 1000 Genomes Evolutionary Extraction Pipeline

**Study:** Metabolic Gating and the Evolution of Human Cognitive Plasticity: A Comparative Genomic Analysis
**Author:** Bryan A. Krantz

## Overview

This repository contains the programmatic extraction pipeline utilized to map the evolutionary trajectory of the "Vanguard" cognitive and metabolic network. 

To determine the evolutionary constraint, archaic introgression, and balancing selection pressures acting upon the loci identified in our GWAS meta-analyses, this Python script systematically queries the 1000 Genomes Project (Phase 3) API. It mathematically isolates the ancestral minor allele frequencies (MAF) for specific Schizophrenia and Scientific Creativity risk alleles across distinct global super-populations.

## Methodology

The pipeline `split_hcar_archaic_miner_robust.py` executes the following workflow:

1. **Locus Ingestion:** Reads the extracted top-tier summary statistics (`.tsv` files) for the ten core Vanguard loci (e.g., *HCAR1/2*, *CACNA1C*, *FOXP2*, *PDHB*).
2. **API Interrogation:** Pings the Ensembl REST API (`https://rest.ensembl.org/variation/human/`) for each highly significant index SNP to retrieve the global phase 3 population data.
3. **Allelic Harmonization:** Automatically resolves strand flips between the GWAS input (A/T/C/G) and the Ensembl reference genome to ensure the correct risk allele frequency is captured.
4. **Super-Population Extraction:** Isolates the exact allelic frequencies for the African (AFR), European (EUR), and East Asian (EAS) super-populations.
5. **Evolutionary Flagging:** Automatically tags "Smoking Gun" variants exhibiting extreme population divergence:
    * *Archaic Introgression Signatures:* High European frequency (>20%) with near-zero African frequency (<5%), indicating post-migration selection or Neanderthal/Denisovan introgression.
    * *Ancestral Baseline Signatures:* High African frequency (>80%) with degraded Eurasian frequency, indicating the original paleolithic hardware blueprint.

## Requirements

The script requires standard Python 3 and the following packages:
* `pandas`
* `requests`

## Outputs

Execution of the script generates `Split_HCAR_Archaic_Divergence.tsv`. This matrix serves as the direct mathematical input for plotting via archaic_divergence_plotter.py.
