## Supplemental Data 7 for "Metabolic Gating and the Evolution of Human Cognitive Plasticity: A Comparative Genomic Analysis": split_hcar_archaic_miner_robust.py

```
import pandas as pd
import requests
import time
import os

# The exact files you generated in Phase 1
FILES = {
    "5' HCAR2 (BHB Cooling Switch)": "HCAR_5prime_cluster_top_SNPs.tsv",
    "3' HCAR1 (Lactate Brake)": "HCAR_3prime_cluster_top_SNPs.tsv",
    'CACNA1C (Calcium Grid)': 'Symbol_CACNA1C_hits.tsv',
    'GRIN2A (Glutamate)': 'Symbol_GRIN2A_hits.tsv',
    'CACNB2 (The Calcium Channel Scaffold)': 'Symbol_CACNB2_hits.tsv',
    'SNAP91 (The Synaptic Vesicle Driver)': 'Symbol_SNAP91_hits.tsv',
    'FOXP2 (The Linguistic/Symbolic Coordinator)': 'Symbol_FOXP2_hits.tsv',
    'BDNF (The Miracle-Gro of the Brain)': 'Symbol_BDNF_hits.tsv',
    'PDHB (The Glycolytic Bottleneck)': 'Metabolism_PDHB_Door_hits.tsv',
    'DRD2 (Dopamine Receptor D2)': 'Symbol_DRD2_hits.tsv'
}

def get_1000g_frequencies(rsid, risk_allele):
    """Pings the Ensembl REST API for 1000 Genomes population frequencies."""
    server = "https://rest.ensembl.org"
    ext = f"/variation/human/{rsid}?pops=1"
    
    # INCREASED TO 5 ATTEMPTS WITH EXPONENTIAL BACKOFF
    for attempt in range(5):
        try:
            # Increased timeout to 10 seconds for laggy connections
            response = requests.get(server + ext, headers={"Content-Type": "application/json"}, timeout=10)
            
            if response.status_code == 429:
                time.sleep(2 * (attempt + 1)) # Back off if rate-limited
                continue
                
            # CATCH SERVER CRASHES (500, 502, 503, 504)
            if response.status_code in [500, 502, 503, 504]:
                time.sleep(3 * (attempt + 1)) # Server is choking, wait longer
                continue
            
            if response.status_code == 400:
                return None, None, "Retired rsID"
                
            if not response.ok:
                return None, None, f"HTTP {response.status_code}"
                
            data = response.json()
            pops = data.get('populations', [])
            
            # Temporary storage for population frequencies by allele
            pop_data = {'EUR': {}, 'AFR': {}, 'EAS': {}}
            pop_map = {
                '1000GENOMES:phase_3:EUR': 'EUR',
                '1000GENOMES:phase_3:AFR': 'AFR',
                '1000GENOMES:phase_3:EAS': 'EAS'
            }
            
            for p in pops:
                p_name = p.get('population')
                if p_name in pop_map:
                    region = pop_map[p_name]
                    allele = p.get('allele')
                    freq = p.get('frequency', 0.0)
                    pop_data[region][allele] = freq
            
            # Resolve the correct frequency for our specific SCZ Risk Allele
            comp = {'A': 'T', 'T': 'A', 'C': 'G', 'G': 'C'}
            comp_allele = comp.get(risk_allele, risk_allele)
            
            freqs = {'EUR': 0.0, 'AFR': 0.0, 'EAS': 0.0}
            matched_allele = None
            
            for region in ['EUR', 'AFR', 'EAS']:
                if risk_allele in pop_data[region]:
                    freqs[region] = pop_data[region][risk_allele]
                    matched_allele = risk_allele
                elif comp_allele in pop_data[region]:
                    freqs[region] = pop_data[region][comp_allele]
                    matched_allele = comp_allele + " (Flip)"
                    
            if matched_allele is None:
                return None, None, "Allele Not Found in 1000G"
                
            return freqs, matched_allele, "Success"
            
        except requests.exceptions.RequestException:
            time.sleep(2 * (attempt + 1)) # Network drop, wait and retry
            continue
            
    return None, None, "Server/Network Timeout (Max Retries Hit)"

def run_split_sweep():
    print("--- Running Split Evolutionary Sweep on Vanguard Network ---")
    results = []
    
    for locus_name, file_path in FILES.items():
        if not os.path.exists(file_path):
            print(f"⚠️ Missing {file_path}. Skipping.")
            continue
            
        df = pd.read_csv(file_path, sep='\t')
        
        # Grab the top 50 most significant SNPs
        top_snps = df.sort_values(by='PVAL').head(50)
        
        print(f"\nScanning: {locus_name} (Testing {len(top_snps)} variants)")
        print("-" * 50)
        
        successful_pings = 0
        for _, row in top_snps.iterrows():
            rsid = row['ID']
            risk_allele = row['A1']
            
            freqs, matched, status = get_1000g_frequencies(rsid, risk_allele)
            
            if freqs:
                print(f"[{rsid}] Risk: {risk_allele} | Matched: {matched} | EUR: {freqs['EUR']:.2f}, AFR: {freqs['AFR']:.2f}")
                successful_pings += 1
                results.append({
                    'Locus': locus_name,
                    'rsID': rsid,
                    'Risk_Allele': risk_allele,
                    'P_Value': row['PVAL'],
                    'EUR_Freq': freqs['EUR'],
                    'AFR_Freq': freqs['AFR'],
                    'EAS_Freq': freqs['EAS']
                })
            else:
                print(f"[{rsid}] ⚠️ Skipped: {status}")
                
            # OPTIONAL: Stop after 15 successful hits per gene to save time. 
            # (Increase this number if you want the script to run longer and dig deeper!)
            if successful_pings >= 15:
                break
                
            time.sleep(0.2) # Be nice to Ensembl servers
            
    if results:
        res_df = pd.DataFrame(results)
        output_file = "Split_HCAR_Archaic_Divergence.tsv"
        res_df.to_csv(output_file, sep='\t', index=False)
        print(f"\n✅ Sweep Complete! Clean data saved to: {output_file}")
        
        # Check for the "Neanderthal" Smoking Gun (High Europe, Zero Africa)
        archaic_guns = res_df[(res_df['AFR_Freq'] < 0.05) & (res_df['EUR_Freq'] > 0.20)]
        print(f"\n🔥 FOUND {len(archaic_guns)} 'ARCHAIC INTROGRESSION' VARIANTS (EUR > 20%, AFR < 5%):\n")
        if not archaic_guns.empty:
            print(archaic_guns[['Locus', 'rsID', 'EUR_Freq', 'AFR_Freq']].to_string(index=False))

        # Check for the "Ancestral Out-of-Africa" Signature (High Africa, Low Europe)
        ancestral_guns = res_df[(res_df['AFR_Freq'] > 0.80) & (res_df['EUR_Freq'] < 0.40)]
        print(f"\n🌍 FOUND {len(ancestral_guns)} 'ANCESTRAL BASELINE' VARIANTS (AFR > 80%, EUR < 40%):\n")
        if not ancestral_guns.empty:
            print(ancestral_guns[['Locus', 'rsID', 'EUR_Freq', 'AFR_Freq']].to_string(index=False))

if __name__ == "__main__":
    run_split_sweep()
```
