## Supplementary figures and images for "Metabolic Gating and the Evolution of Human Cognitive Plasticity: A Comparative Genomic Analysis"

### Figure_Hypothesis_Archaic_Divergence.png

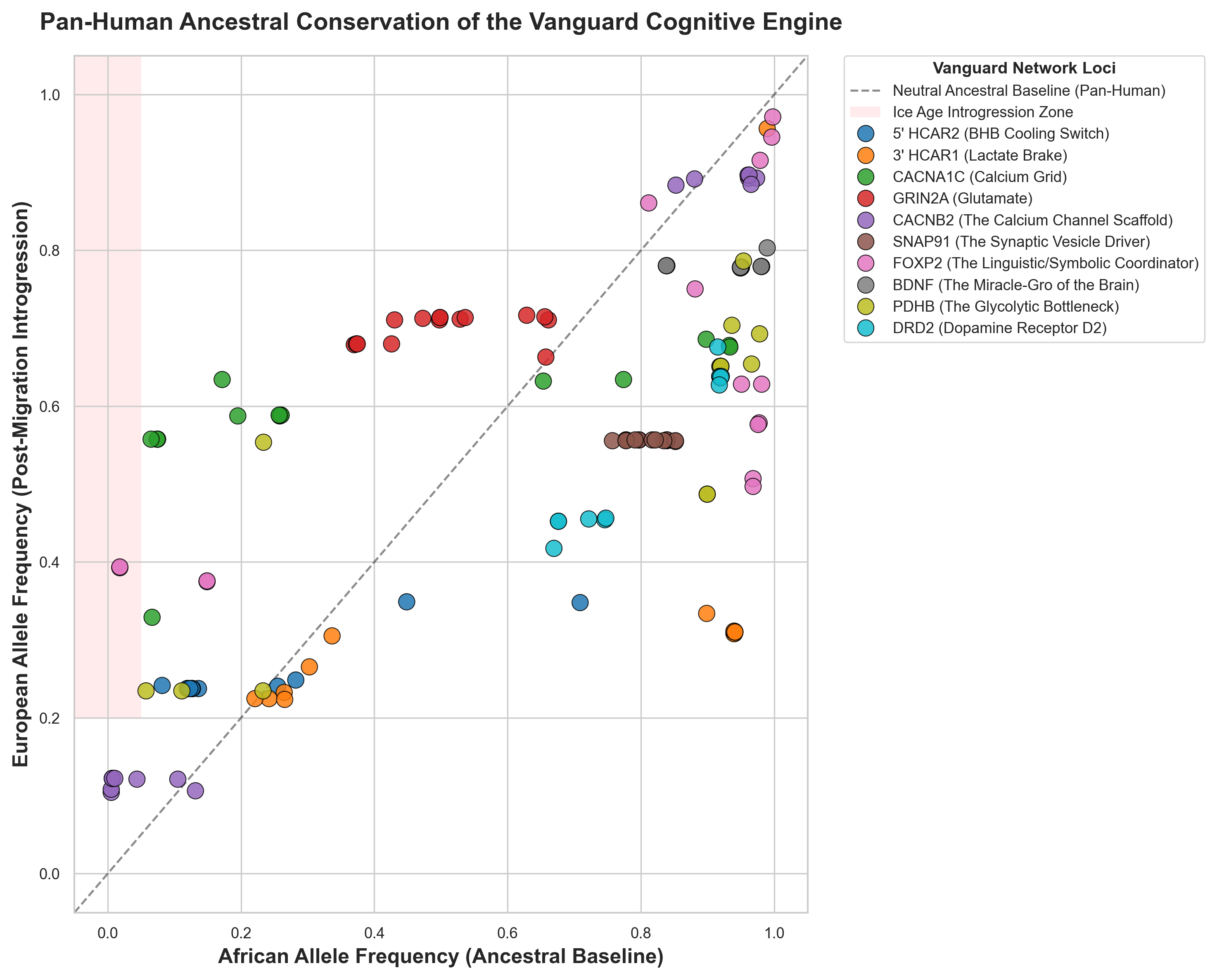
